# Recognition Mechanism of Serotonin by a G-Quadruplex-Duplex Hybrid Aptamer

**DOI:** 10.64898/2026.06.26.734732

**Authors:** Guohua Xu, Chen Wang, Meiting Kang, Jiawen Chen, Junhan Wei, Qiang Zhao, Maili Liu, Conggang Li

## Abstract

Serotonin is a key neurotransmitter, and aptamer-based tools using the 44-nt Apt44 have been successfully developed for its *in vitro* and *in vivo* detection. Nevertheless, the structural basis of recognition by this aptamer remains unclear. Here we report high-resolution NMR structures of Apt38, a 6-nt truncated variant in the third loop of Apt44, in free and serotonin-bound states. Both structures reveal a two-layered antiparallel chair-type G-quadruplex core with three edgewise loops and a terminal duplex, forming a G-quadruplex–duplex hybrid structure. Serotonin binds at the G-quadruplex–duplex junction, stabilized by stacking, electrostatic attraction, hydrogen bonding, and hydrophobic contacts. Apt38 is preorganized for binding, whereas the longer third loop of Apt44 introduces conformational dynamics into the G-quadruplex scaffold, which enables a pronounced binding-triggered conformational switch in PBS buffer, explaining its sensing mechanism. Our work reveals the recognition and sensing mechanism of the serotonin aptamer and provides a framework for aptamer design in serotonin biosensing.

## Introduction

Serotonin (5-hydroxytryptamine, 5HT) is a key neurotransmitter that regulates diverse functions of the central and peripheral nervous systems, including mood, sleep, appetite, memory, learning and cognition. Dysregulation of serotonin signaling is implicated in a wide range of neuropsychiatric disorders, including depression, anxiety, schizophrenia, and autism spectrum disorder. Consequently, the ability to detect serotonin with high sensitivity and specificity, as well as to monitor its dynamic fluctuations in real time, particularly in complex *in vivo* biological environments, is essential for both basic neuroscience research and clinical diagnostics.^[1-10]^

Aptamers are single-stranded oligonucleotides isolated via the Systematic Evolution of Ligands by Exponential Enrichment (SELEX) process, capable of recognizing a broad spectrum of targets—from metal ions and small molecules to proteins and even whole cells—with high affinity and specificity. Their advantages, including ease of synthesis, flexible chemical modification, and low immunogenicity, render them highly attractive for applications in biosensing, disease diagnosis, therapeutics and drug delivery.^[11-20]^ These features make aptamers ideal candidates for developing selective serotonin sensors.

In 2018, Nakatsuka et al. selected a 44-mer guanine (G)-rich DNA aptamer with high affinity (*K*_d_ ≈30 nM) for serotonin (Figure 1a-b).^[21]^ Despite the small size of serotonin and its structural resemblance to other neurotransmitters such as dopamine and norepinephrine, this aptamer exhibits remarkable specificity, positioning it as a promising molecular recognition tool. Taking advantage of the significant conformational change induced by serotonin binding, the aptamer-based field-effect transistor (FET) sensors were first successfully developed and achieved the sensitive detection of serotonin under physiological ionic strength. Subsequently, this aptamer has been employed in various sensing platforms, including field-effect transistor, fluorescent and electrochemical assays,^[22-35]^ showing great potential in broad applications.

**Figure 1.**
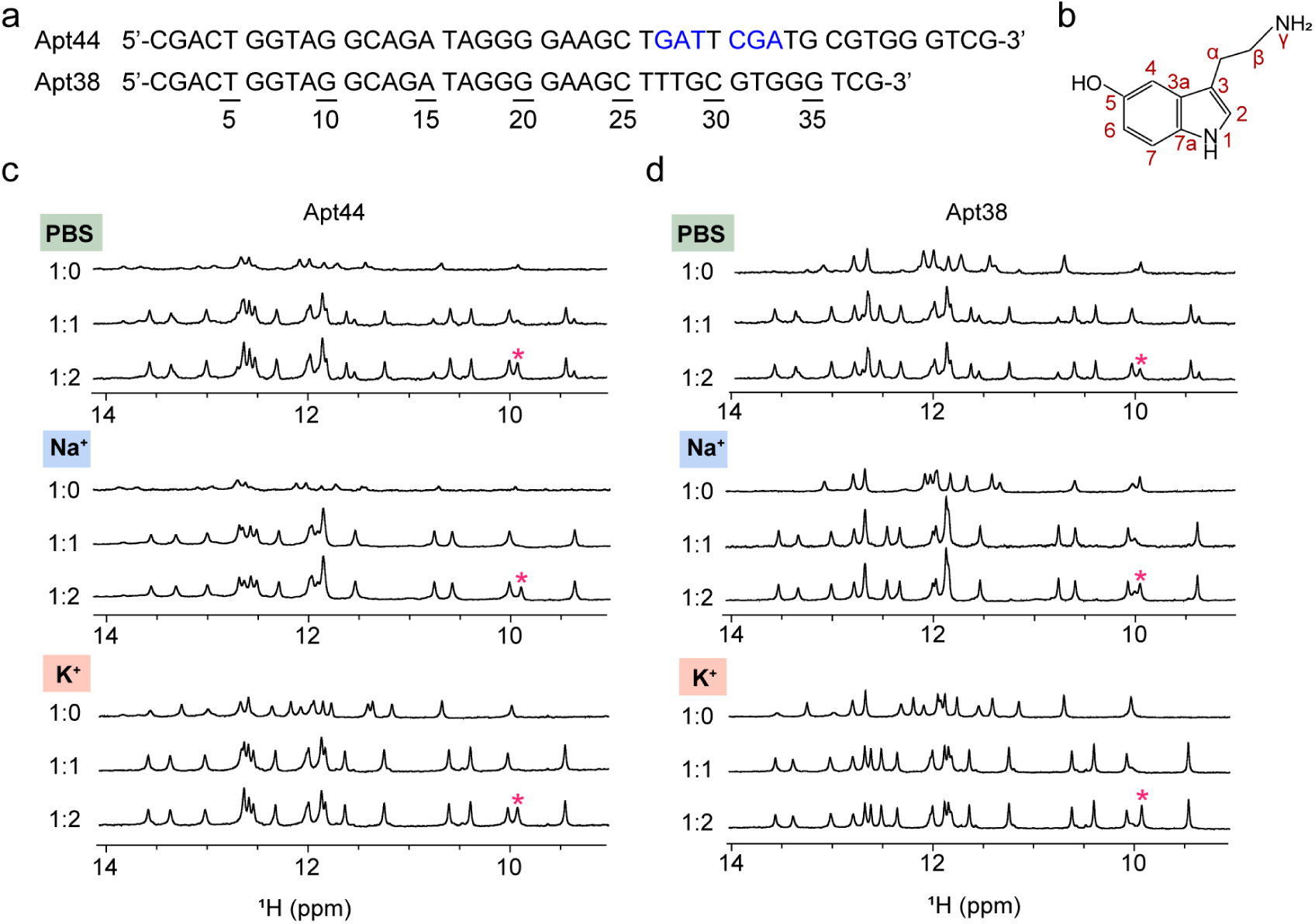
^1^H NMR spectra of the serotonin aptamer. (a) Sequence of 44-mer serotonin aptamer (Apt44) and its truncated variant, Apt38. (b) Chemical structure of serotonin. (c) Imino region of ^1^H NMR spectra of Apt44 in the absence and presence of serotonin under three buffer conditions (PBS, Na⁺ buffer, and K⁺ buffer) at 298 K. (d) Imino region of ^1^H NMR spectra of Apt38 in the absence and presence of serotonin under the same conditions as in (c). The asterisk (*) denotes the resonance from free serotonin.

Despite its excellent performance in sensor applications of this aptamer, the molecular mechanism underlying serotonin recognition remains poorly understood at the detailed structural level. Circular dichroism (CD) spectroscopy indicated serotonin induced the aptamer to shift away from predominant duplex to an antiparallel G-quadruplex.^[21]^ Recent efforts combining molecular dynamics simulations, molecular docking, and single-molecule field-effect transistor measurements have begun to elucidate potential binding sites, serotonin-induced conformational rearrangements.^[27, 35-37]^ Nevertheless, the absence of an experimentally determined high-resolution structure of the serotonin–aptamer complex leaves fundamental questions unresolved: What conformational changes does the aptamer undergo before and after serotonin binding? What is the architecture of the binding pocket? Which key residues and interactions govern its high affinity and specificity for serotonin in complex physiological environments where structurally similar molecules coexist?

Here, we report the high-resolution NMR structures of the serotonin aptamer in its free and bound states. The structures reveal a G-quadruplex-duplex hybrid architecture with the binding pocket at the junction of G-quadruplex and duplex—a binding mode distinct from prior computational predictions. These findings provide a structural basis for high-affinity serotonin recognition by its aptamer and are expected to facilitate the rational design of serotonin aptamers for neurochemical sensing applications and drug discovery, as well as to contribute to the broader understanding of nucleic acid–small molecule recognition.

## Results and Discussion

### The Structural Property of Serotonin Aptamer Investigated by NMR and CD

The imino region signals in ¹H NMR spectra of nucleic acids provide key structural information: signals at 10–12 ppm typically correspond to G-quadruplexes (arising from guanine H1), while signals at 12–14 ppm indicate duplex regions (arising from guanine H1 in G•C base pairs and thymine H3 in A•T base pairs). In addition, the linewidth of these signals can be used to assess conformational homogeneity and regularity, with sharper lines indicating a more homogeneous and well-ordered structure. To investigate the structural properties of serotonin aptamer Apt44, whose high affinity was confirmed by isothermal titration calorimetry (ITC) (Figure S1), we firstly acquired its ^1^H NMR spectra in phosphate-buffered saline (PBS) supplemented with 2 mM MgCl₂ (Figure 1c) – the same buffer in which Apt44 was selected by SELEX. The presence of signal peaks in the 10–14 ppm region, both in the absence and presence of serotonin, indicates that Apt44 contains both G-quadruplex and duplex structural elements. However, it adopts a better-defined structure upon serotonin binding than in its free state. The ¹H NMR spectrum remained unchanged upon addition of two equivalents of serotonin compared to that with one equivalent, suggesting a 1:1 binding stoichiometry, in agreement with the ITC result (Figure S1).

To determine whether the G-quadruplex is stabilized by K⁺ or Na⁺, we further acquired ¹H NMR spectra of Apt44 in buffers containing either K⁺ or Na⁺, each with MgCl₂ (Figure 1c). The comparison of NMR spectra of Apt44 in PBS, K⁺ and Na⁺ buffers shows that under the PBS conditions used (which contain high Na⁺ and low K⁺ concentrations), Apt44 adopts a G-quadruplex conformation stabilized by Na⁺ in the absence of serotonin, whereas it primarily adopts a G-quadruplex conformation stabilized by K⁺ in the presence of serotonin. Additionally, in K⁺ buffer, the aptamer adopts a well-defined G-quadruplex and duplex structure even in the absence of serotonin, indicating that K⁺ can stabilize the aptamer conformation in both the free and bound states more effectively than Na⁺.

We also performed circular dichroism (CD) analysis on the aptamer in PBS, K⁺ buffer, and Na⁺ buffer (Figure S2). The results are consistent with the NMR findings, supporting that the aptamer forms a structure comprising an antiparallel G-quadruplex in the presence of serotonin. This is evidenced by the appearance of a negative band at 260 nm and positive bands at 240 nm and 290 nm upon serotonin binding in PBS, K⁺ buffer, and Na⁺ buffer. Additionally, the nearly identical CD profiles before and after serotonin addition in K⁺ buffer indicate the well-defined antiparallel G-quadruplex has formed for the free aptamer in K⁺ buffer.

### Truncation of the Aptamer to Obtain a Construct Suitable for NMR Structural Determination

Considering the high-quality ¹H NMR spectra of the aptamer in K⁺ buffer for both the free and bound states, we proceeded to acquire 2D ^1^H-^1^H COSY, TOCSY, and NOESY spectra of Apt44 under K⁺ condition, to obtain complete ^1^H signal assignments and ultimately determine the high-resolution structure. However, during spectral analysis, we found that completing assignment of all ¹H resonances was challenging using these data alone due to spectral overlap. In order to simplify spectra and improve spectra quality, we decided to generate a truncated variant of Apt44 based on the limited structural information available for the aptamer. Given the presence of an antiparallel G-quadruplex conformation indicated by CD and ^1^H NMR spectra, and the formation of Watson–Crick base pairs between 5’-CGAC and GTCG-3’ revealed by the partial assignments of Apt44, we hypothesized several possible structural models for Apt44 (Figure S3). In these models, a long loop region of 17-19 nucleotides was present. Therefore, we attempted to shorten this loop. Compared to the imino region of the ¹H NMR spectrum of Apt44, a truncated variant (Apt38), in which six nucleotides were deleted from the loop, exhibited highly similar spectra both in the absence and presence of serotonin (Figure 1a and 1d). ITC further confirmed that Apt38 retains a binding affinity comparable to that of Apt44 (Figure S1). Based on these findings, we proceeded with the shorter Apt38 construct for subsequent NMR structural studies.

### Proton Resonance Assignments of Free and Serotonin-bound Apt38

Following the approach used for Apt44, we acquired 1D and 2D NMR spectra of Apt38 in K⁺ buffer and completed the assignment of the ¹H NMR signals both in the absence and presence of serotonin (Table S1 and S2). For the Apt38–serotonin complex, H5–H6 and CH₃–H6 cross-peaks from cytosine and thymine bases were unambiguously observed in TOCSY and COSY spectra, facilitating the identification of base types for H8/H6 resonances in the NOESY spectra. The well-resolved NOESY spectra enabled us to trace H8/H6–H1′ sequential connectivities for the majority of nucleotides, with the exception of the A13–G19 region, where these connectivities were partially missing (Figure 2b). H8/H6–H2′/H2′′/H3′ NOE connectivities were used to further confirm the assignments. Guanine and thymidine imino (H1 and H3) protons were assigned using 1D ¹⁵N-edited HMQC spectra on 3% ¹⁵N site-specifically labeled samples (Figure 2a and S4). Thymidine H3 assignments were further confirmed by NOE correlations to their respective methyl groups. Adenosine H2 protons were assigned using 1D ¹³C-edited HMQC spectra on 5% ¹³C site-specifically labeled samples (Figure S5).

**Figure 2.**
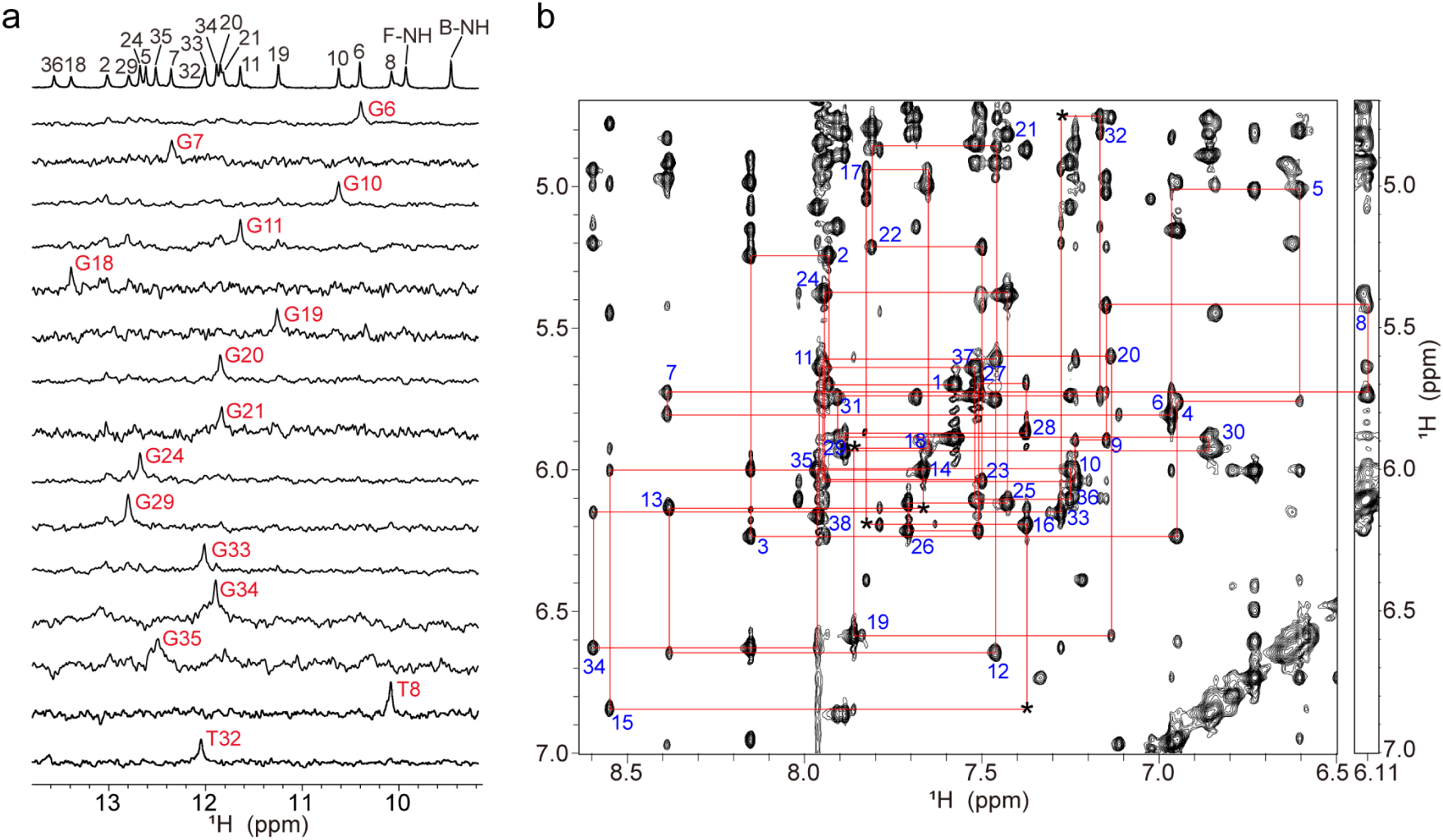
Proton NMR assignments of Serotonin-bound Apt38 in K⁺ buffer. (a) The imino region of 1D ^1^H NMR spectra of Apt38-serotonin complex. Imino protons were assigned by 1D ^15^N-edited HMQC spectra using 3% site-specifically ^15^N-labeled samples. F-NH and B-NH represent the NH protons of free and bound serotonin, respectively. (b) NOESY spectrum of Apt38-serotonin complex (300 ms mixing time), showing the H8/6-H1′ sequential connectivities. Intraresidue H8/6-H1′ cross-peaks are labeled with residue numbers. Missing connectivities are marked with asterisks.

After identifying the proton signals for the majority of bases, the H8/H6 resonances of A13–G19 region where sequential H8/H6–H1′ connectivities were partially missing, were further assigned as follows: (i) the H8 resonances of several adenosine bases were identified by exploiting the characteristic NOEs of adenosine, in which H1′ exhibits intraresidual NOEs with both H8 and H2 (the latter being weak); (ii) the H6 resonance of T16 was identified based on the only unassigned CH₃–H6 signal; and (iii) the remaining sequential connectivities, including G14H1′–A15H8 and A17H1′–G18H8, were used to identify the H8 resonances of guanine bases. The proton signals of bound serotonin in the NMR spectra were assigned based on detailed analysis of the serotonin chemical structure and by utilizing exchange peaks between free and bound serotonin. Based on the proton resonance assignments of the complex, we proceeded to assign the free Apt38 using a similar approach (Figure S6 and S7). The assignment of the free aptamer was relatively straightforward, facilitated by the spectral similarity between the free and serotonin-bound forms.

### The Solution Structure of the Apt38

By analyzing NOEs between imino (H1 and H3) and other protons based on the resonance assignments (Figure 3a-b), the overall folding topology of the free Apt38 and Apt38-serotonin complex was determined. For both states, characteristic imino-H8 proton cyclic NOE connectivity patterns around G-tetrads revealed a two-layered, chair-type antiparallel G-quadruplex core, formed by G6•G11•G19•G34 and G7•G33•G20•G10 tetrads (Figure 3c-d). The glycosidic conformations of the guanosine residues within these tetrads are *syn*•*anti*•*syn*•*anti* for the first tetrad and *anti*•*syn*•*anti*•*syn* for the second tetrad, respectively, as reflected by H8-H1’ NOE cross-peak intensities (Figure 2b). Within the G-quadruplex core, the two G-tracts G6-G7 and G19-G20 run in one direction, while the other two G-tracts, G10-G11 and G33-G34, run in the opposite direction (Figure 3d). The G-quadruplex structure features three edgewise loops: a 2-nt loop (T8-A9), a 7-nt loop (C12-G18) and a 12-nt loop (G21-T32). Additionally, the bases C1-G2-A3-C4 and G35-T36-C37-G38 form duplex through canonical Watson-Crick base pairing (G38•C1, G2•C37, A3•T36, G35•C4 pairs). This duplex formation is supported by the strong NOEs between the imino proton of G bases and the amino protons of C base, between the imino proton of T base and the H2 proton of A base, as well as by the NOEs observed between adjacent Watson-Crick base pairs. Taken together, these data indicate that both free and serotonin-bound Apt38 adopt a structure comprising an antiparallel chair-type G-quadruplex and a duplex.

**Figure 3.**
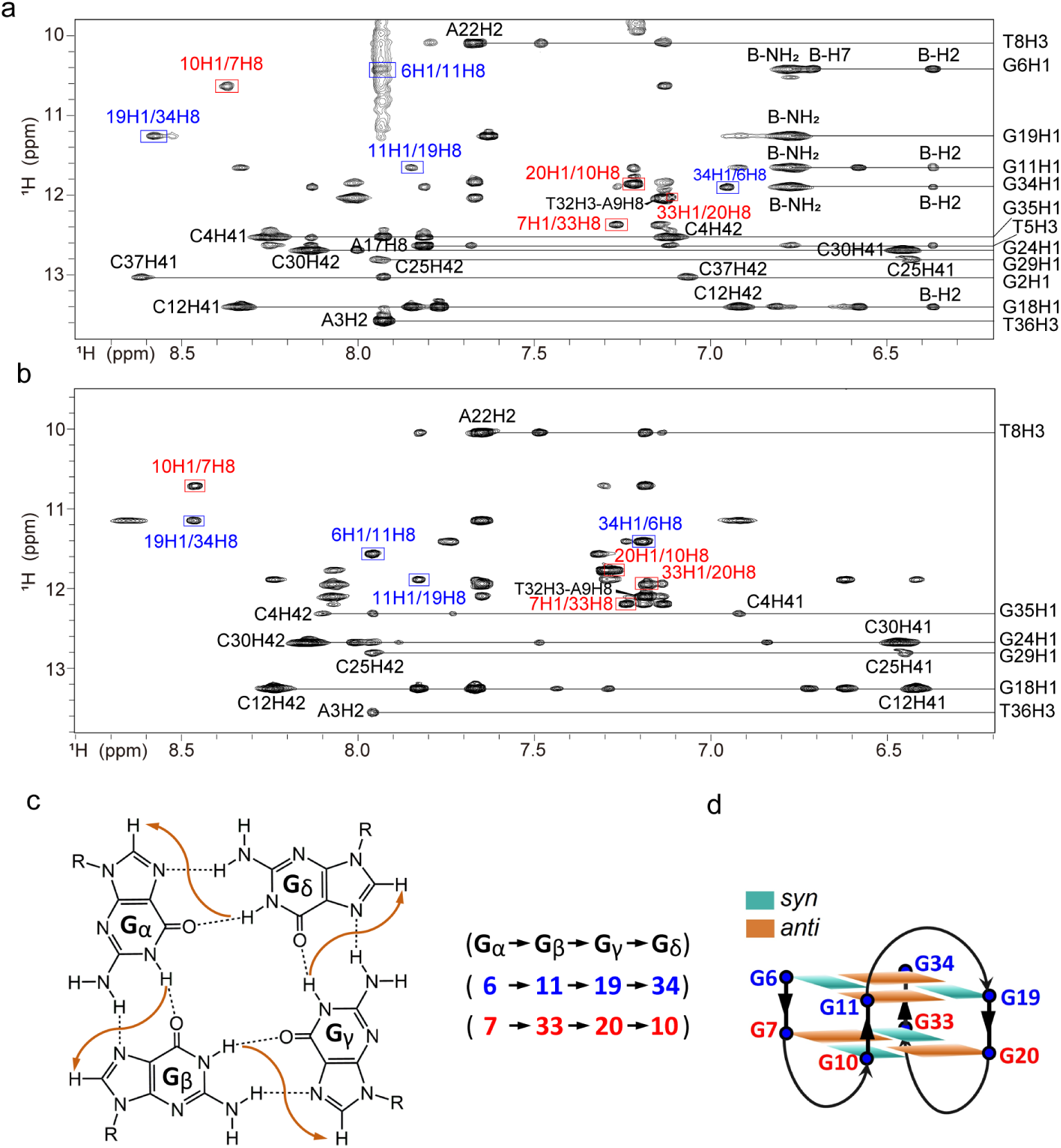
NOESY spectra of free Apt38 and Apt38-serotonin complex in K^+^ buffer (90%H_2_O/10%D_2_O) at 298 K. (a) Expanded NOESY spectrum of Apt38-serotonin complex, correlating NOEs between imino protons and other protons. The cross-peaks showing characteristic imino-H8 proton cyclic NOE connectivity around the two G-tetrads, G6•G11•G19•G34 and G7•G33•G20•G10, are framed in blue and red, respectively. B indicates the bound serotonin, and B-NH_2_, B-H7, and B-H2 represent proton resonances of the bound serotonin in crosspeaks with the aptamer. (b) Expanded NOESY spectrum of free Apt38 under the same conditions, with the corresponding cyclic imino–H8 proton NOE connectivity patterns for the two G-tetrads framed in blue and red, respectively. (c) Schematic representation of G-tetrads observed in the NOESY spectrum. Characteristic imino-H8 protons NOE connectivity pattern around the G-tetrad plane is shown in orange row. (d) Schematic illustration of G-quadruplex core for free Apt38 and Apt38-serotonin complex, where *anti* and *syn* guanines within the G-tetrads are colored orange and cyan, respectively.

Building on these topological insights, the high-resolution structures of free Apt38 and its complex with serotonin were calculated using Xplor-NIH, based on NMR restraints including intramolecular and intermolecular NOEs, hydrogen bonds, and dihedral angles (Tables S3 and S4). The lowest-energy refined structures are shown in Figure 4. Both the free and bound structures are well defined, with average pairwise heavy-atom RMSD values of 1.34 ± 0.26 Å and 1.47 ± 0.47 Å, respectively, excluding the flexible loop residues 14–16 and 26–28 (Table S4). The representative structures of free Apt38 and the complex are highly similar, with an all-atom RMSD of 2.39 Å.

**Figure 4.**
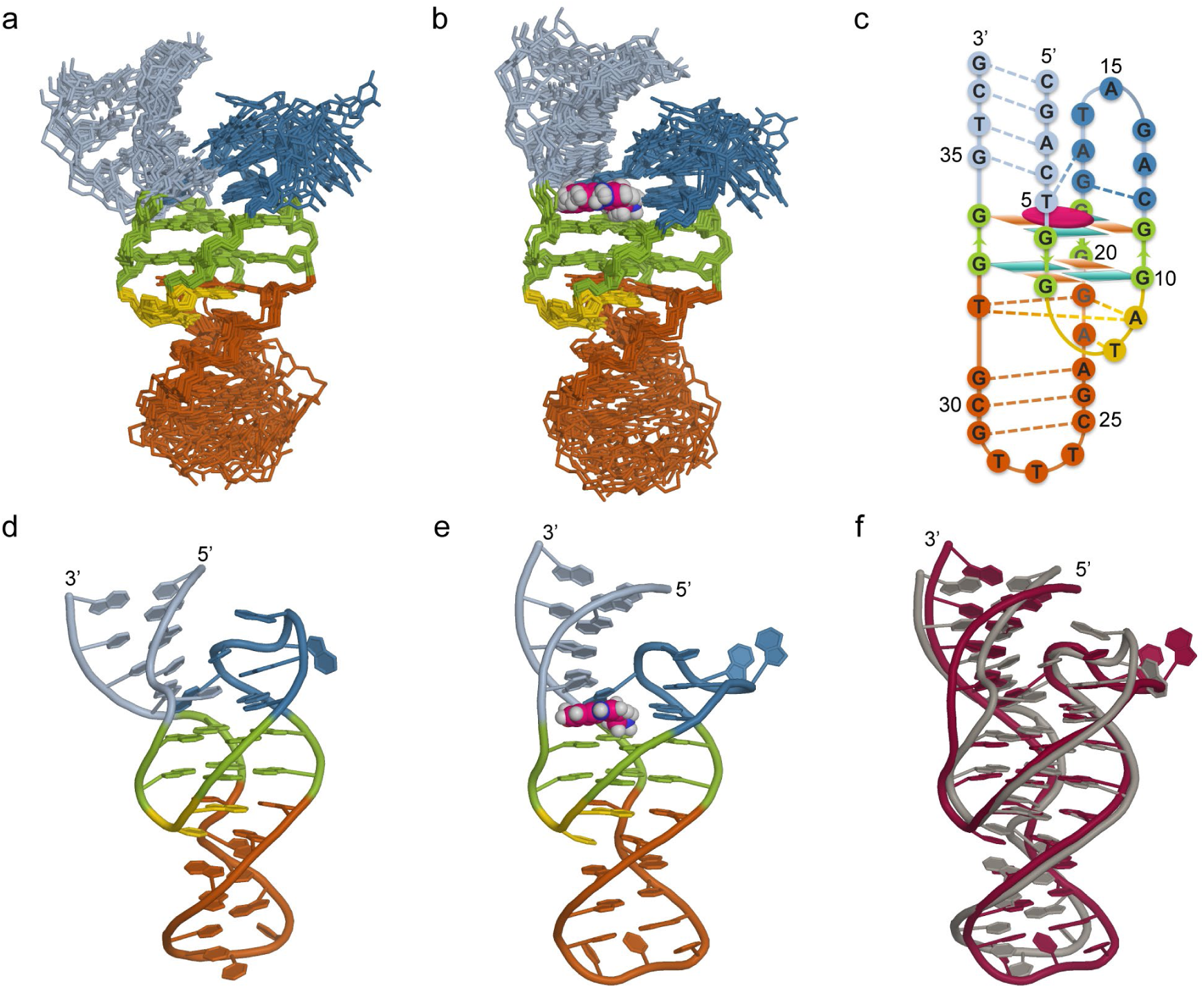
Solution structures of free Apt38 and the Apt38-serotonin complex. (a, b) Ten superimposed refined structures of free Apt38 (a) and Apt38-serotonin complex (b). The structures were aligned except for residues G14–T16 and T26–T30 in the loops. Serotonin is shown as a sphere model. (c) Schematic illustration of Apt38-serotonin complex, where *anti* and *syn* guanines within the G-tetrads are colored orange and cyan, respectively. Serotonin is shown in purple ellipse. (d-e) Cartoon representation of a refined structure of free Apt38 (d) and Apt38-serotonin complex (e). (f) Structural overlap of free Apt38 (gray) and Apt38-serotonin complex (purple).

The structure shows that the G-quadruplex construct has three edgewise loops, i.e., the first 2-nt loop T8-A9, the second 7-nt loop C12-G18 and the third 12-nt loop G21-T32. The stacking interactions between the two G-tetrads, together with these inter- and intra-loop base pairings, collectively stabilize the G-quadruplex structure (Figure 5a). The base pairings, both inter- and intra-loop, include the A9•G21•T32 triple and the mismatched T8•A22 pair formed between the first loop and the third loop; the G18•C12 Watson-Crick pairing formed within the second loop; and the G24•C30 and G29•C25 Watson–Crick pairs, along with the mismatched G31•A23 pair formed within the third loop (Figure 5b).

**Figure 5.**
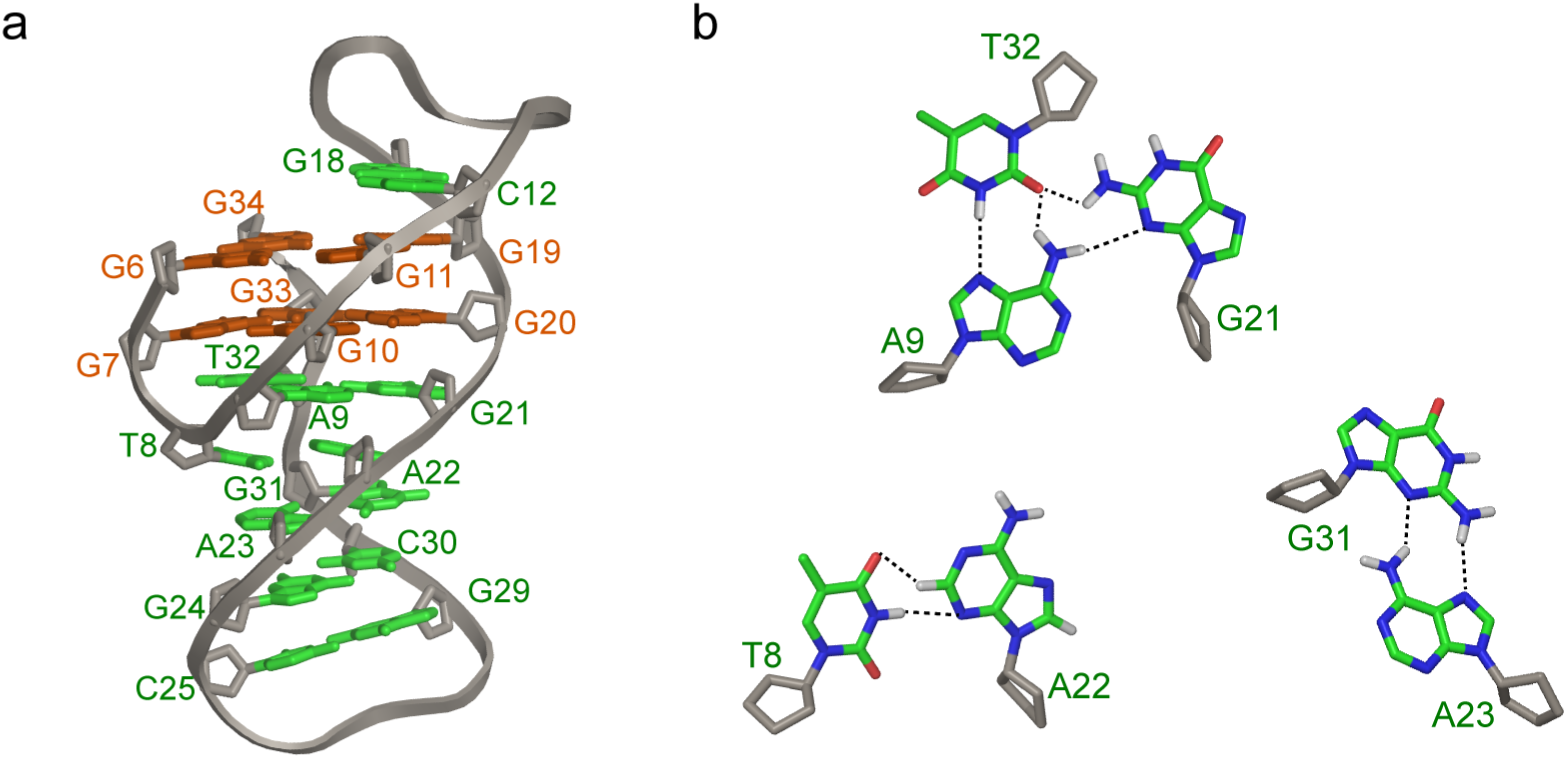
The stacking interactions between the two G-tetrads, together with these inter- and intra-loop base pairings, collectively stabilize the G-quadruplex structure. (a) Side view showing the stacking between the two G-tetrad planes (G6•G11•G19•G34 and G7•G33•G20•G10), together with base pairings both between and within loops. These include the A9•G21•T32 triple, the mismatched T8•A22 pair connecting the 2-nt loop (T8-A9) and the third 12-nt loop (G21-T32), the Watson-Crick pairs G24•C30 and G29•C25 along with the mismatched G31•A23 pair within the third 12-nt loop, and the Watson-Crick pair G18•C12 within the second 7-nt loop. (b) Hydrogen-bonding interactions in the A9•G21•T32 triple, the T8•A22 pair, and the G31•A23 pair. Sugars are colored gray. The oxygen, nitrogen, carbon and hydrogen atoms for bases are depicted in red, blue, green and light gray, respectively.

### Binding Pocket Architecture

Structural analysis revealed that the binding site is composed of a G-tetrad plane (G6•G11•G19•G34) together with two base pairs, G18•C12 and A17•T5. Serotonin is sandwiched between the G-tetrad plane and the A17•T5 base pair, with the side sterically confined by G18·C12 (Figure 6a-c). Notably, the terminal amino group of serotonin is positioned directly above the center of the G-tetrad plane. Clearly, π–π stacking interactions between the aromatic ring of serotonin and both the A17•T5 base pair and the G-tetrad plane, together with hydrophobic contacts involving the aromatic protons of serotonin with the T5 sugar ring and the G6-H5’/H5’’ atoms, constitute the primary driving forces for specific recognition by the aptamer. Furthermore, the terminal amino group of serotonin has a p*K*_a_ of approximately 10 and is therefore predominantly protonated as –NH_3_ ^+^ under the neutral buffer conditions used in this study. Consequently, a significant electrostatic attraction exists between this protonated amino group and the negatively charged G-tetrad plane. In addition, the –NH_3_ ^+^ group can also form hydrogen bonds with the O6 atoms of the four guanines in the G-tetrad, as well as with the N7 and O6 atoms of G18 (Figure 6d). Collectively, π–π stacking, hydrophobic effects, hydrogen bonding, and electrostatic interactions synergistically contribute to the high-affinity binding of serotonin to the G-quadruplex aptamer.

**Figure 6.**
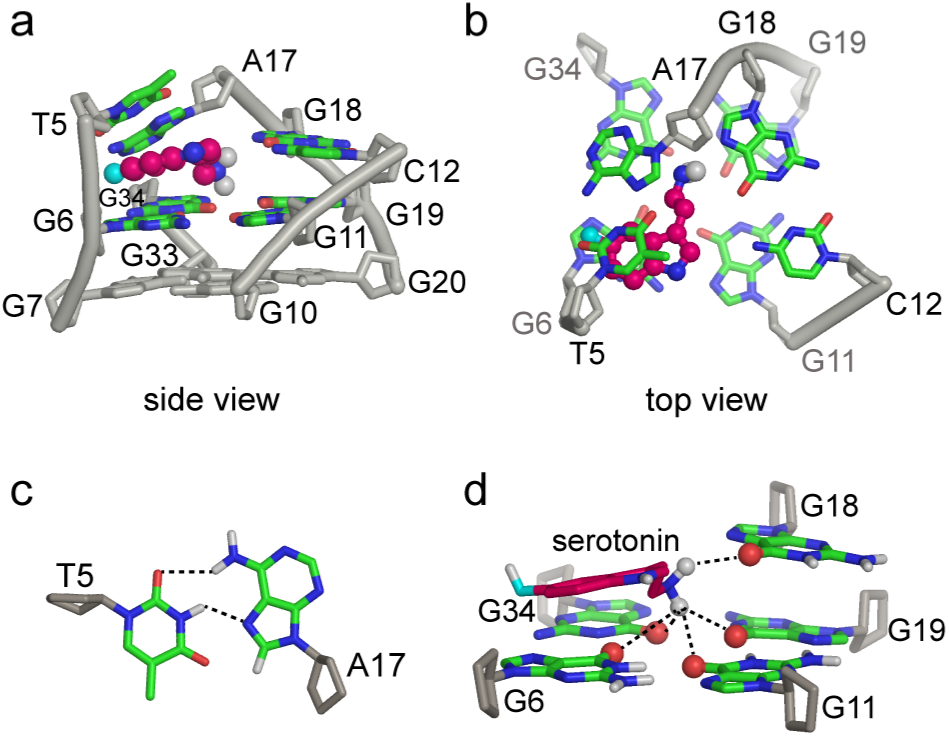
Structural details of the serotonin-binding site in a representative Apt38–serotonin complex structure. (a) Side view of the serotonin-binding pocket. Serotonin is shown in a ball-and-stick model. (b) Side view of the serotonin-binding pocket showing the aromatic ring of serotonin stacking between T5 of the A17•T5 base pair and G6 of the G-tetrad. (c) Hydrogen-bonding interactions of the A17•T5 base pair. (d) Side view showing the hydrogen bonds between the amino group of serotonin and the O6 atoms of the four guanines in the G-tetrad, as well as the O6 atom of G18. Hydrogen bonds are shown as dashed lines.

In addition, comparison of the free and bound aptamer structures revealed only minor conformational differences. The most notable difference between the two states lies in the phosphate backbone segment from T5 to G6 within the binding site (Figure 4f). In the free state, this segment exhibits a more bent conformation, whereas upon serotonin binding, it becomes straighter and more extended, consistent with the ligand inserting into the binding pocket and filling the space to create a more compact and fully occupied binding site. Together, these observations support a conformational selection mechanism.

### Ligand-Induced Structural Stabilization of Apt38 Detected by NMR

The resolved structures of free and bound Apt38 showed only minor conformational differences, and CD spectroscopy revealed almost no overall conformational change upon serotonin binding, as reflected by nearly superimposable spectra before and after ligand addition (Figure S2). However, 1D ^1^H NMR spectra of the imino proton region provided critical information on local dynamics. The NH signals corresponding to T36 and G2 in the duplex region (base pairs T36•A3 and G2•C37) exhibited distinct linewidth changes upon serotonin binding (Figure 1d). In K⁺-containing buffer, these signals were visible but significantly broadened in the free state; upon binding, they became markedly sharper. In Na⁺-containing buffer, the same NH signals were completely invisible in the free state—likely due to enhanced dynamics or rapid exchange—but emerged as sharp peaks after serotonin binding. Notably, the imino signal of T5, which forms the T5•A17 base pair that bridges the G-quadruplex core and the duplex region, was also absent in the free state under both K⁺ and Na⁺ conditions but appeared as a sharp peak upon serotonin binding. These observations collectively indicate that, in the free state, the duplex region and its junction with the G-quadruplex core are relatively dynamic and heterogeneous, rendering several imino signals invisible or significantly broadening. Upon serotonin binding, the ligand inserts into the binding site and establishes a continuous, tightly packed stacking network that connects the duplex region with the G-quadruplex core. This effectively stabilizes the duplex region and rigidifies the overall structure. Together, NMR reveals pronounced ligand-induced suppression of local dynamics and stabilization of overall structure.

### Critical Base Pairs Governing Serotonin Binding

To verify the correctness of the resolved structure and identify the base pairs critical for serotonin binding, we performed systematic base mutation experiments. Because Apt38 and Apt44 share nearly identical core folding (as demonstrated by the structural comparison in the next section), we carried out systematic mutation experiments primarily using Apt44, where a more comprehensive mutant library was available (Table S5, S6 and S7). Within the binding site, introducing G18A, G18I, or C12T mutations in Apt44 completely abolished serotonin binding, while the A17T mutation or substitution of T5 with A/C/G drastically reduced binding affinity, indicating that the G18•C12 and A17•T5 base pairs within the binding site are essential for high-affinity interaction. In addition, several loop-region pairs that are distant from the binding site but stabilize the G-quadruplex structure were also found to be critical for affinity, including the G21•A9•T32 triplet (where T32 corresponds to T38 in Apt44), A22•T8, A23•G31, G24•C30, and G29•C25 (G29, C30, G31 correspond to G35, C36, and G37 in Apt44, respectively). ITC results showed that introducing A9C, G21C, or T8A mutations in Apt38, as well as A9T, T8C, G21A, A22T, T38C, A23T, C36T or G37T mutations in Apt44, completely disrupted serotonin binding, and the A23C mutation in Apt44 also caused a sharp decrease in affinity (*K*_d_ ≈ 625 ± 199 nM), confirming the conservation and importance of the A9•G21•T32 triplet and the A23•G31 and A22•T8 pairs. These mutational results validate the resolved structure and demonstrate that high-affinity serotonin binding requires both the direct recognition elements within the binding site and the structural stability of the G-quadruplex scaffold.

Beyond the binding site and loop regions, the duplex region also contributes to high-affinity binding. Within the duplex region, the C4•G35 base pair in Apt38 (corresponding to C4•G41 in Apt44) forms stacking interactions with the T5•A17 base pair from the binding site. The functional importance of this C4•G41 pairing was assessed by introducing three mutations in Apt44 that replace the original C4•G41 pair with G4•C41, A4•T41, and T4•A41, respectively. All these mutations led to reduced binding affinity, with *K*_d_ values of 321 nM, 415 nM, and 1750 nM, respectively. These results confirm that the C4•G41 base pair is critical for high-affinity binding—a finding that can be rationalized by the tightly stacking interaction between C4•G35 and T5•A17 observed in the resolved Apt38 structure. In addition, ITC results revealed that retaining four base pairs in the duplex is the minimum required for high affinity, whereas further extension did not significantly improve affinity.

### Loop Length Modulates G-Quadruplex Conformational Dynamics: A Comparative Study of Apt38 and Apt44

Having established that both the binding site and distant base pairs (including loop-region pairs and the duplex pair) are critical for high-affinity recognition, we next examined the functional role of the loop regions themselves. In both the second and third G-quadruplex loops, there are flexible turn regions. Using Apt44, we found that these regions are not conserved for target recognition, as demonstrated by a series of mutation experiments (Table S5 and S7). For the second loop (nucleotides A13–T16 in Apt44), mutations such as A13T, G14A, A15T, and T16C did not disrupt aptamer affinity. Similarly, single-base mutations in the third loop, including C25T, T26C, G27A, A28T, T29G, T30G, C31T, G32A, A33T, and T34C, did not significantly change affinity.

However, the third loop length does affect structural stability (Table S6 and S7). For instance, the truncated variant Apt38, in which six nucleotides are deleted from the flexible turn region of the third G-quadruplex loop, is more stable than the parent Apt44. Specifically, Apt38 folds into a well-ordered, stable G-quadruplex structure in both K⁺ and Na⁺ buffers in the free state (Figures 1 and S2), and exhibits higher affinity for the target. In contrast, free Apt44 forms a well-folded G-quadruplex only in K⁺ buffer, but not in Na⁺ buffer, where it exhibits much broader NMR linewidths and lacks the characteristic G-quadruplex CD signal. These observations confirm that the loop-enhanced dynamics of Apt44 significantly affect the structural homogeneity and stability of the G-quadruplex scaffold, representing the major distinction between Apt38 and Apt44. This distinction becomes smaller after binding to serotonin, owing to the formation of continuous and tight stacking interactions.

The difference in conformational rigidity and dynamics between Apt38 and Apt44 also directly determines their respective advantages. Apt38, with its higher structural stability, facilitates the acquisition of higher-quality NMR spectra and thus benefits structural determination, but it is less suitable for sensor designs that rely on binding-induced conformational changes, as it undergoes little change upon serotonin binding. In contrast, the greater dynamics introduced by the longer loop of Apt44 confers a distinct advantage in sensor applications. For instance, in PBS buffer with high Na⁺ and low K⁺ concentrations, Apt44 undergoes a pronounced conformational transition from an unstable Na⁺-stabilized G-quadruplex to a more stable K⁺-stabilized G-quadruplex upon serotonin binding (Figure 1 and Figure S2). Such a binding-induced conformational switch underpins the widespread use of Apt44 in sensing applications.

### The Binding Analysis of Serotonin Analogs and Aptamer

To clarify the contributions of key functional groups to binding, we performed NMR and ITC studies of serotonin analogs binding to Apt44/Apt38 (Figure S8 and Table S8). It was found that substitutions at the 5-hydroxy position of serotonin (e.g., tryptamine, 5-chlorotryptamine, 5-methyltryptamine, 5-methoxytryptamine) even enhanced binding affinity toward the aptamer. NMR showed that these substitutions primarily affected the chemical shifts of the spatially adjacent T5 and G6 bases. Structural analysis revealed that the 5-hydroxy site is laterally open with no base obstruction, and thus introduction of different groups does not impair binding. We propose that the main driving force for the enhanced binding of all these analogs relative to serotonin is enhanced hydrophobic interaction, because replacing the polar hydroxyl with more hydrophobic groups (H, Cl, CH₃, or OCH₃) reduces the desolvation penalty within the binding pocket.

In contrast, analogs with a modified terminal amino group of serotonin, such as tryptophol and 5-hydroxy-L-tryptophan, completely lost binding ability. Based on the NMR structure, this loss can be explained by the distinct properties of each analog. Tryptophol, which replaces the amino group with a neutral hydroxyl, cannot form electrostatic interactions with the G-quadruplex plane. Although 5-Hydroxy-L-tryptophan retains the amino group, the α-carboxyl introduces steric hindrance and the molecule exists as a zwitterion (net charge zero) at physiological pH, preventing effective approach of the amino group to the G-quadruplex plane. These results suggest that electrostatic attraction between the protonated amino group and the G-quadruplex plane is essential for binding. These findings also have important implications for using Apt44 as a serotonin sensing tool. Naturally occurring 5-substituted analogs, particularly tryptamine, may cause significant cross-interference due to their strong binding affinity to the aptamer. In contrast, metabolites with a modified terminal amino group (e.g., tryptophol, 5-hydroxy-L-tryptophan) do not interfere.

### NMR Structure versus Computational Binding Models

Previous studies have proposed potential binding models of the serotonin aptamer Apt44, as well as the conformational changes of the aptamer upon serotonin binding, using molecular dynamics simulations and/or molecular docking analyses^[27, 36-37]^. However, the NMR solution structure obtained in this work reveals that the actual binding mode and conformational change mechanism are not consistent with those previous computational models. This discrepancy may primarily arise from the current limitations of common nucleic acid three-dimensional modeling methods, such as 3DRNA, which still struggle to accurately construct complex G-quadruplex-duplex topologies and recapitulate the native folding backbone of the aptamer, thereby potentially leading to deviations from the true binding mode. This suggests that, at present, obtaining accurate binding models for nucleic acid–small molecule interactions involving complex topologies remains highly dependent on direct experimental determination. Concurrently, the development of high-precision modeling algorithms, specialized force fields, and advanced conformational sampling methods for complex nucleic acid architectures represents a critical direction for future progress in the field. The synergistic iteration between experimental structures and theoretical simulations will jointly promote the long-term advancement of mechanistic studies on nucleic acid aptamer recognition and rational design.

### The G-quadruplex–Duplex Junction: A Conserved Structural Hotspot for Small-Molecule Recognition

NMR structural analysis indicates that the serotonin aptamer adopts a well-defined G-quadruplex-duplex hybrid architecture in solution, with the ligand-binding pocket located at the junction between the G-quadruplex core and the duplex region. Notably, this quadruplex-duplex junction has been previously identified as a high-affinity binding site for various small molecules, including G-quadruplex ligands such as pyridostatin derivatives and Phen-DC3, the classical OTA aptamer, as well as artificially designed ligands reported in the literature^[38-44]^. For instance, certain G-quadruplex ligands (e.g., pyridostatin derivatives and Phen-DC3) intercalate directly at the interface between the G-quadruplex and a coaxially stacked duplex. In contrast, serotonin stacks onto the G-quadruplex plane but not at the G-quadruplex-duplex interface, forming its binding site through the G-quadruplex plane in cooperation with flexible residues at the junction. OTA, on the other hand, does not directly contact the G-quadruplex plane; instead, it relies on the duplex region together with the flexible junction residues to create its binding pocket. Despite these differences in binding modes, all observations suggest that the G-quadruplex-duplex junction may serve as a recurrent hotspot for small-molecule recognition.

We further analyzed the physicochemical properties of these small molecules and found that those capable of direct stacking with the G-quadruplex plane are generally positively charged or contain an extended aromatic ring system that facilitates stacking interactions (e.g., pyridostatin derivatives and Phen-DC3). The G-quadruplex plane carries a strong negative electrostatic potential; under physiological conditions, such small molecules are favored to interact with this negatively charged surface through stacking, electrostatic attraction, or hydrogen bonding. Conversely, negatively charged molecules such as OTA experience electrostatic repulsion from the G-quadruplex plane, making direct stacking unfavorable. Instead, OTA appears to achieve specific recognition through halogen bonding, hydrophobic interactions, and stacking with duplex bases. Notably, despite their distinct electrostatic properties, all of these small molecules form functional binding cavities at the G-quadruplex-duplex junction, underscoring the versatility and compatibility of this region in accommodating diverse ligands.

Taken together, these observations may point to a conserved recognition mechanism for aptamers adopting such a structural architecture. The G-quadruplex and duplex serve as rigid scaffolds that provide spatial confinement and stacking interfaces, while the junction between them can function as a versatile binding platform. Depending on the ligand’s structure and electrostatic properties, this platform may accommodate binding either through direct intercalation at the rigid G-quadruplex–duplex interface or via engagement with flexible residues at the junction to form an adaptive pocket. This mechanistic insight provides a structural basis for the rational design of ligands targeting this hotspot and for the development of modular aptamer architectures.

## Conclusion

In this study, we determined the high-resolution NMR solution structures of the serotonin aptamer Apt38 in both its free and serotonin-bound states. The structures reveal a two-layered antiparallel chair-type G-quadruplex core with three edgewise loops and a terminal duplex for both states. Serotonin binds in a pocket formed by a G-tetrad plane, the A17•T5 base pair, and the G18•C12 base pair, stabilized by stacking, electrostatic attraction, hydrogen bonding, and hydrophobic contacts. Apt38 undergoes only minor conformational changes upon serotonin binding, as its free state already adopts a stable structure resembling the bound state. In contrast, the longer third loop of the parent Apt44 introduces conformational dynamics into the G-quadruplex scaffold, which enables a pronounced conformational switch upon serotonin binding in PBS buffer. Such a binding-induced conformational switch underpins the widespread use of Apt44 in sensing applications. Collectively, our work establishes a clear structure–dynamics–function relationship for serotonin aptamers and offers a rational framework for aptamer design.

## Supporting information

Experimental section and Figure S1-S8 Table S1-S8

## Supporting Information

NMR spectra, CD spectra, ITC titration curves, Sequence and ITC-determined dissociation constant of aptamers, Proton chemical shifts, Statistics of the computed structures, Intermolecular NOEs data, including Figures S1–S8 and Tables S1–S8.

## Accession Codes

The coordinates and chemical shifts for the free Apt38 aptamer and its serotonin complex have been deposited in the Protein Data Bank and the Biological Magnetic Resonance Data Bank (BMRB). Accession codes are as follows: free aptamer (PDB: 22QK, BMRB: 36824) and serotonin complex (PDB: 21Ll, BMRB: 36818).

## Acknowledgments

This work was financially supported by National Natural Science Foundation of China grants (22474148, 22407127); the Strategic Priority Research Program of the Chinese Academy of Sciences (XDB0540000, YSBR-068 and XDB0750100), National Key R&D Program of China (2024YFA0918802).

## Table of Contents

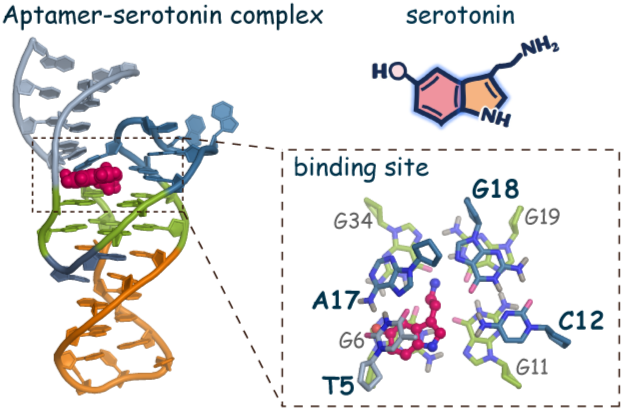

