## Supplementary material for "Recognition Mechanism of Serotonin by a G-Quadruplex-Duplex Hybrid Aptamer": Experimental section and Figure S1-S8 Table S1-S8

#### 1. Sample preparation

Serotonin hydrochloride was purchased from Sigma-Aldrich (Shanghai) Trading Co., Ltd. and dissolved in $D_2O$ to prepare a 50 mM stock solution. Sodium 2,2-dimethyl-2-silapentane-5-sulfonate (DSS) and isotopically labeled phosphoramidites were obtained from Cambridge Isotope Laboratories, Inc. Site-specifically labeled DNA oligonucleotides were chemically synthesized on an ABI 394 DNA/RNA synthesizer and purified by HPLC using a C18 OBD Prep column. Unlabeled DNA oligonucleotides (HPLC-purified) were supplied by Sangon Biotechnology Co., Ltd. (Shanghai, China). The lyophilized DNA samples were dissolved in either $D_2O$ or a 90% $H_2O$/10% $D_2O$ (v/v) buffer, annealed by heating to 95 °C for 7 min followed by slow cooling to room temperature. DNA concentration was determined by measuring the UV absorbance at 260 nm using the corresponding extinction coefficient.

For 1D $^1H$ NMR spectroscopy, the following buffers were used: phosphate-buffered saline (PBS, 137 mM NaCl, 2.7 mM KCl, 10 mM $Na_2HPO_4$, 1.8 mM $KH_2PO_4$, pH 7.4) supplemented with 2 mM $MgCl_2$, $Na^+$ buffer (20 mM $Na_2HPO_4$/$NaH_2PO_4$, 100 mM NaCl, 2 mM $MgCl_2$, pH 7.4), or $K^+$ buffer (20 mM $K_2HPO_4$/$KH_2PO_4$, 100 mM KCl, 2 mM $MgCl_2$, pH 7.4). The final aptamer concentrations in these experiments were 0.05–0.1 mM. For 2D NMR experiments aimed at structural determination, the aptamer concentrations were increased to 1.0–1.5 mM, and $K^+$ buffer supplemented with 20 mM $MgCl_2$ was used to obtain high-quality spectra. The DNA oligonucleotides were dialyzed to remove residual small molecules from HPLC purification. This was performed by first dialyzing against PBS, followed by dialysis against water.

#### 2. NMR spectroscopy

NMR spectra were acquired on Bruker NMR spectrometers operating at 600, 700, and 850 MHz, each equipped with a CryoProbe and at temperatures of 279 K or 298 K, respectively. For structural analysis, 2D $^1H$–$^1H$ TOCSY (mixing time 120 ms), COSY, and $^{13}C$–$^1H$ HSQC spectra were recorded in $D_2O$. 2D $^1H$–$^1H$ NOESY spectra were collected in $D_2O$ and in $H_2O$ containing 10% (v/v) $D_2O$, using jump-and-return (JR) or W5 water suppression, with mixing times of 120 ms and 300 ms. All 2D spectra were acquired with 256-512 complex points in the $t_1$ dimension and 2048 points in the $t_2$ dimension. $^1H$ chemical shifts were referenced to 2,2-dimethylsilapentane-5-sulfonic acid (DSS) at 0 ppm. For the unlabeled DNA aptamer, 2D $^{15}N$–$^1H$ HMQC spectra were obtained with 12 k scans. In the case of site-specific low-enrichment DNA samples (3% $^{15}N$-labeled guanine or thymine, or 5% $^{13}C$-labeled adenosine), 1D $^{15}N$- or $^{13}C$-edited HMQC spectra were acquired to assign the imino protons of guanine and thymine residues, as well as the adenosine H2 protons. All NMR data were processed using Bruker TopSpin software and analyzed with Sparky.[1]

### 3. Structural calculation

The structures of the free aptamer and its complex were calculated using Xplor-NIH 2.47,[2-3] according to the standard protocol[4] as described previously.[5] The calculations incorporated hydrogen-bonding restraints, intramolecular and intermolecular NOE-derived distance restraints, glycosidic dihedral angles restraints, and planarity restraints for base pairs and G-tetrads. Distance restraints were primarily derived from NOESY spectra recorded with a mixing time of 120 ms. From an initial pool of 1000 refined structures, the 10 lowest-energy models were selected for subsequent analysis. All structural analysis and visualization were performed using the PyMOL Molecular Graphics System, Version 3.0.3 (Schrödinger, LLC).

### 4. Isothermal titration calorimetry (ITC) analysis

ITC was applied to determine binding affinities of aptamers (or mutated sequences) to serotonin by using Microcal PEAQ-ITC microcalorimeter (Malvern). Aptamers and serotonin solutions were prepared in the binding buffer (PBS with 2 mM $MgCl_2$). In ITC measurements, 100 μM serotonin in the injection syringe was titrated into 10 μM aptamers or mutated sequences in sample cell at 298 K. After an initial equilibration for 60 s, titrations started with the first 0.4 μL of serotonin solution and 19 successive 2.0 μL of serotonin solution every 100 s with the syringe stirring speed of 750 rpm. The reference power was set as 10 μcal/s. By integrating the heat pulse area of each titration, the ITC titration curves were obtained, and dissociation constant ($K_d$), binding stoichiometry (N), enthalpy change (ΔH), free energy change (ΔG) and entropy change (TΔS) were finally determined.

### 5. Circular dichroism (CD) spectroscopy

The DNA samples used for CD spectroscopy were diluted to 10 μM in $K^+$ buffer, $Na^+$ and PBS buffer with 2 mM $MgCl_2$, respectively. CD spectra of aptamer (Apt38 or Apt44) and aptamer-serotonin complex were recorded using a Chirascan spectropolarimeter (Applied Photophysics Limited. UK) from 220 to 320 nm at room temperature with a cell path length of 0.1 cm, a step size of 1 nm, a dwell time of 0.5 s/nm, and a bandwidth of 1 nm. Each CD spectrum was an average of three measurements.

### Supplementary Figures and Tables

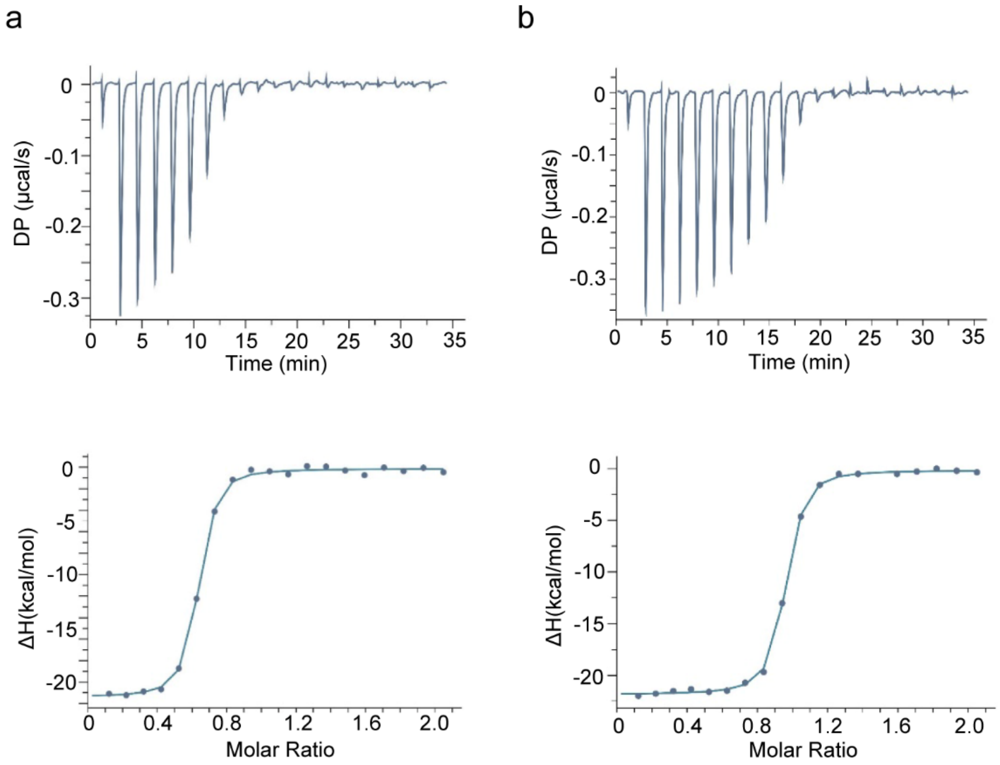


**Figure S1.** ITC titration curves for the binding of aptamers and serotonin. (a) Apt44 ($K_d$ = 37.0 ± 4.3 nM). (b) Apt38 ($K_d$ = 25.4 ± 2.1 nM). Binding buffer: 1×PBS, 2 mM $MgCl_2$, pH 7.4.

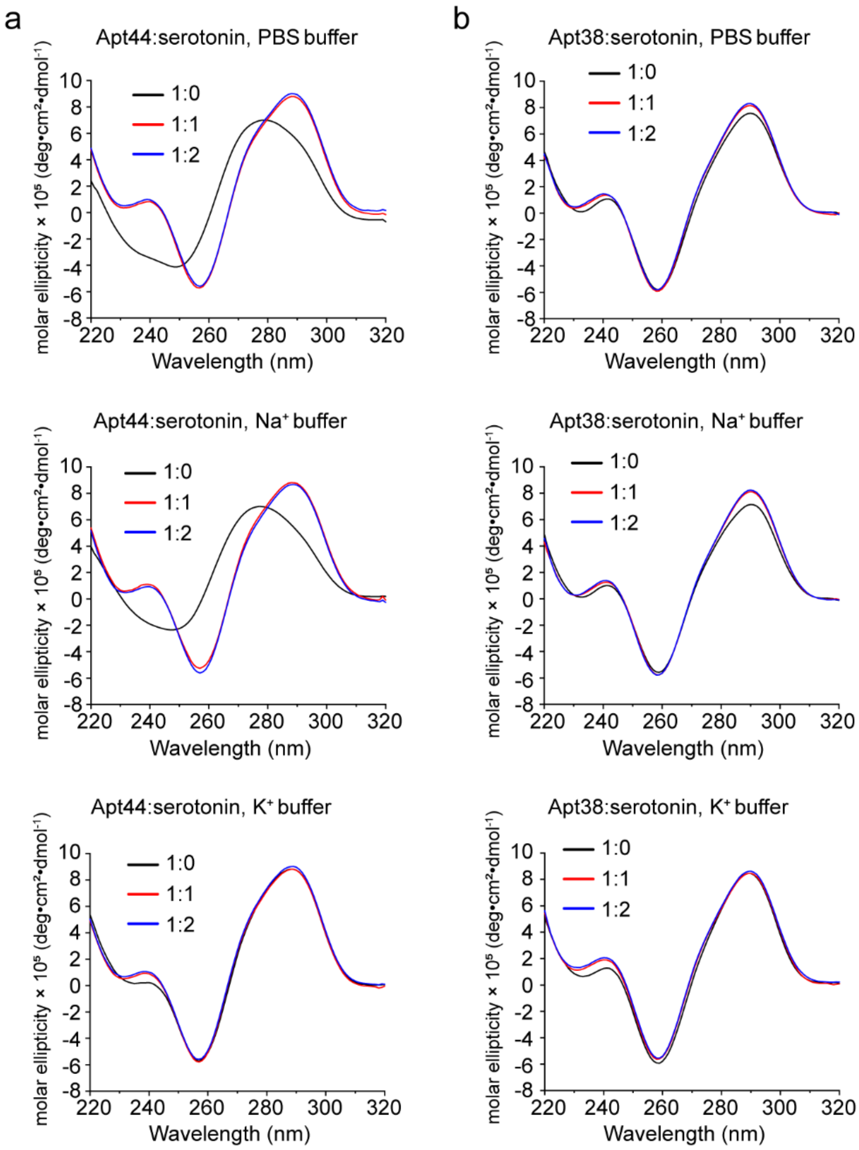


**Figure S2.** CD spectra of Apt44 (a) and Apt38 (b) measured in various buffers containing 2 mM $MgCl_2$, with or without serotonin.

Apt44 5’-CGACT GGTAG GCAGA TAGGG GAAGC TGATT CGATG CGTGG GTCG-3’

5 10 15 20 25 30 35 40

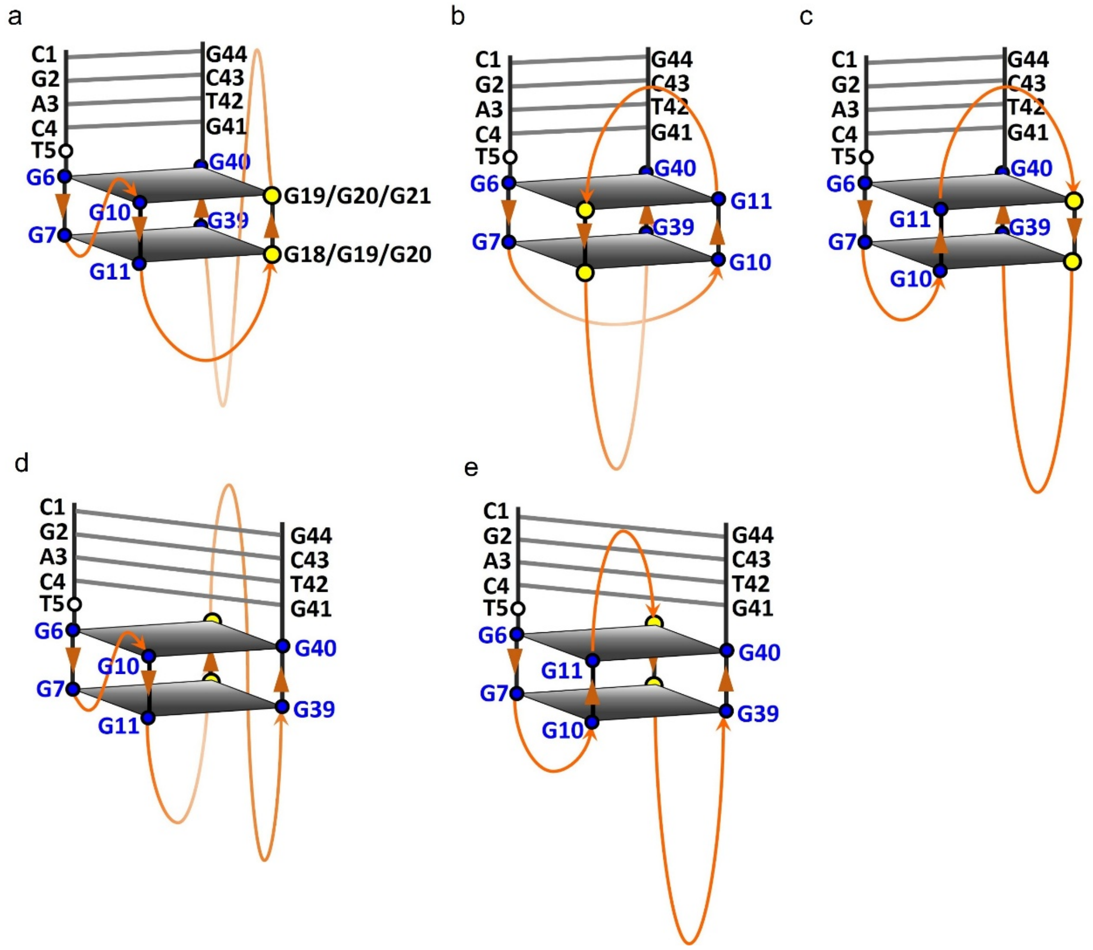


**Figure S3.** Proposed structural models for Apt44 based on the presence of an antiparallel G-quadruplex conformation and the formation of Watson–Crick base pairs between the 5’-C1G2A3C4-3’ and 5’-G41T42C43G44-3’ segments of Apt44. Because the partial G-rich segments involved in G4 formation contain only two consecutive guanines (G6-G7, G10-G11, G39-G40, highlighted in blue), the antiparallel G4 was proposed to consist of two G-quartet layers. Among the four consecutive guanines G18-G21 (highlighted in yellow), which two participated in G4 formation remained unclear, as shown in model a. The same ambiguity applies to G18-G21 in other models and is not annotated.

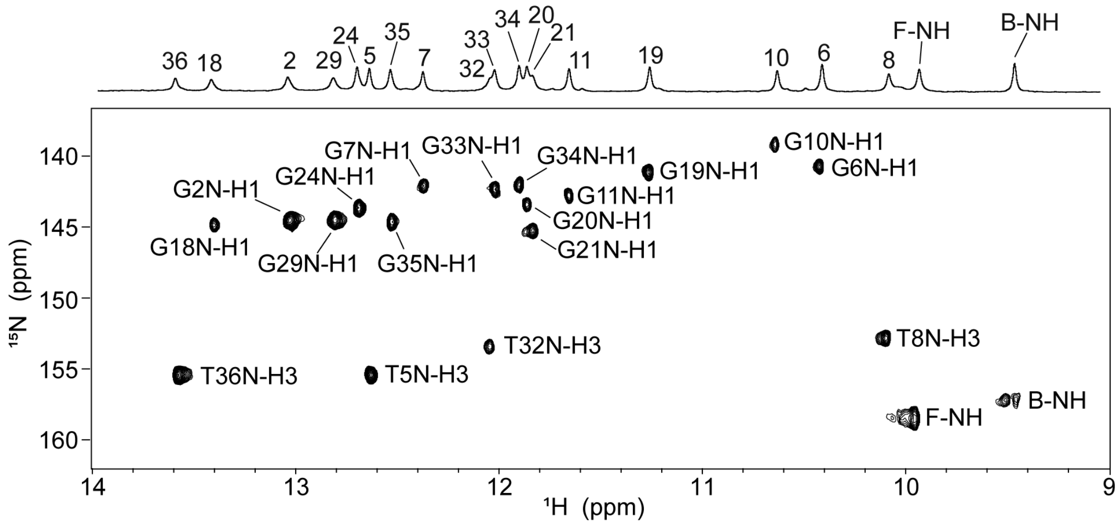


**Figure S4.** 2D $^{15}$N-$^{1}$H HMQC spectra of Apt38-serotonin complex at natural abundance.

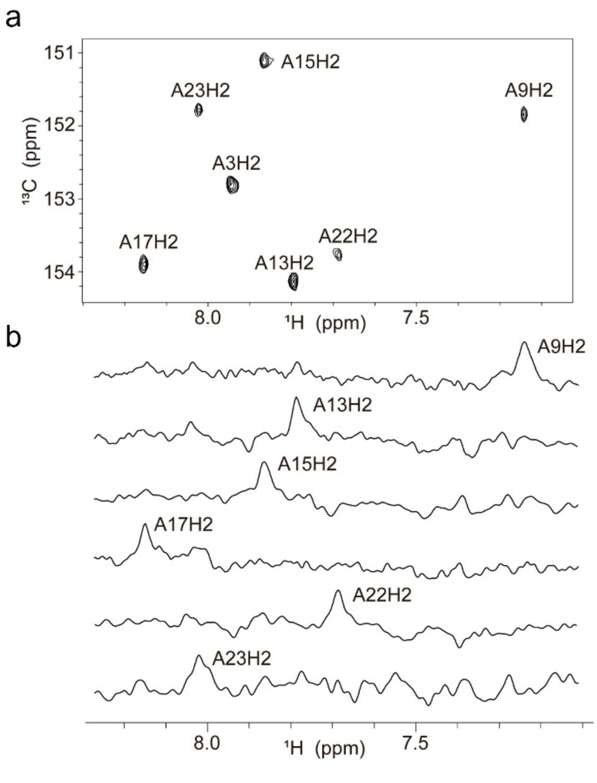


**Figure S5.** The assignment of adenosine H2 protons for Apt38-serotonin complex. (a) 2D $^{13}$C-$^{1}$H HMQC spectra on natural abundance sample. (b) 1D $^{13}$C-edited HMQC spectra on 5% $^{13}$C site-specifically labeled samples.

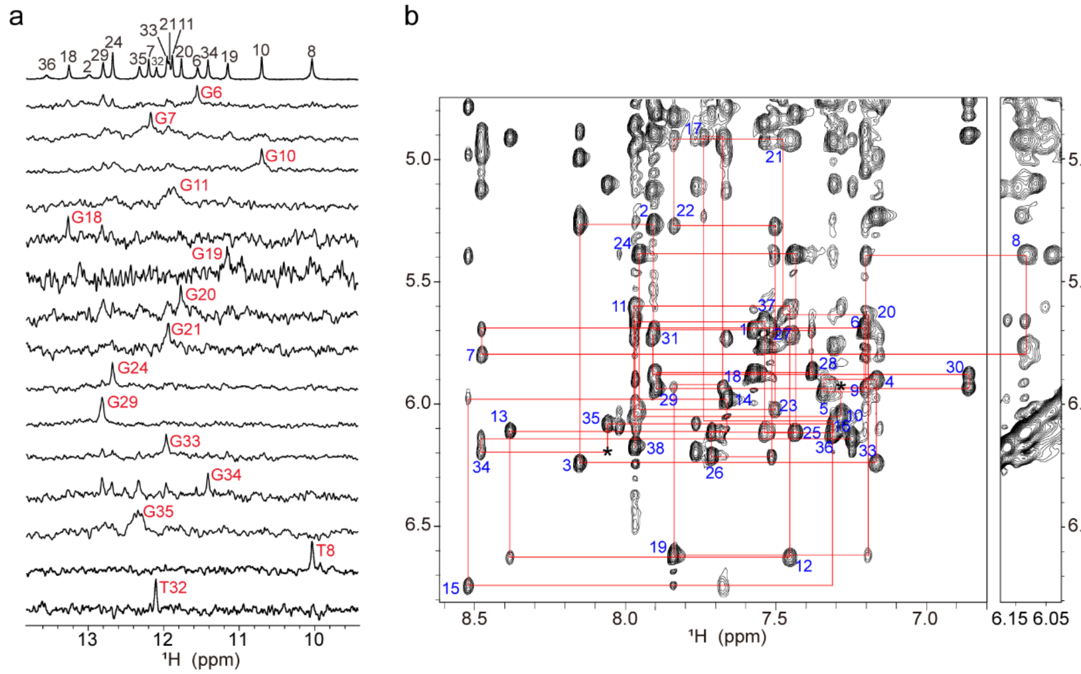


**Figure S6.** Proton NMR assignments of free Apt38 in $K^+$ buffer. (a) The imino region of 1D $^1H$ NMR spectra of free Apt38. Imino protons were assigned via 1D $^{15}N$-edited HMQC spectra using 3% site-specifically $^{15}N$-labeled samples. (b) NOESY spectrum (300 ms mixing time) of free Apt38, showing the H8/6-H1′ sequential connectivities. Intraresidue H8/6-H1′ cross-peaks are labeled with residue numbers. Missing connectivities are marked with asterisks.

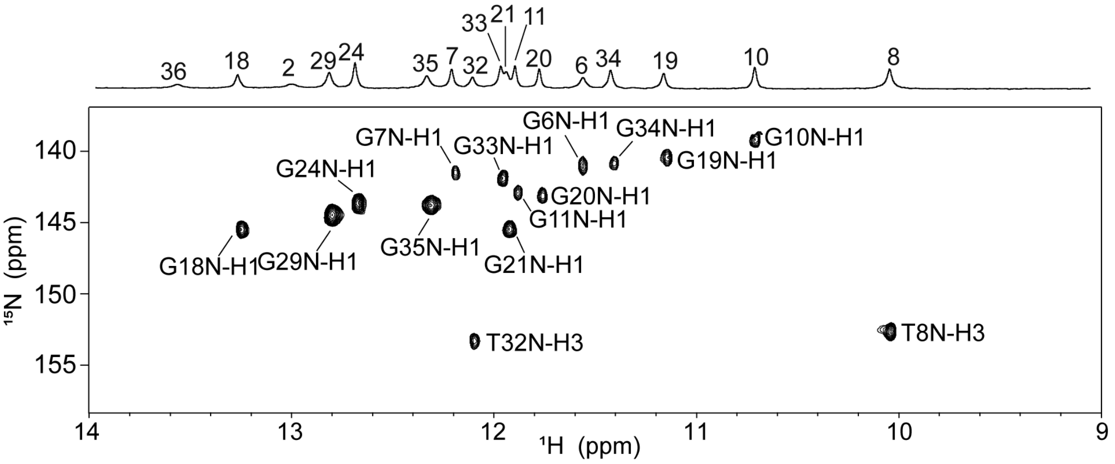


**Figure S7.** 2D $^{15}N$-$^1H$ HMQC spectra of free Apt38 at natural abundance.

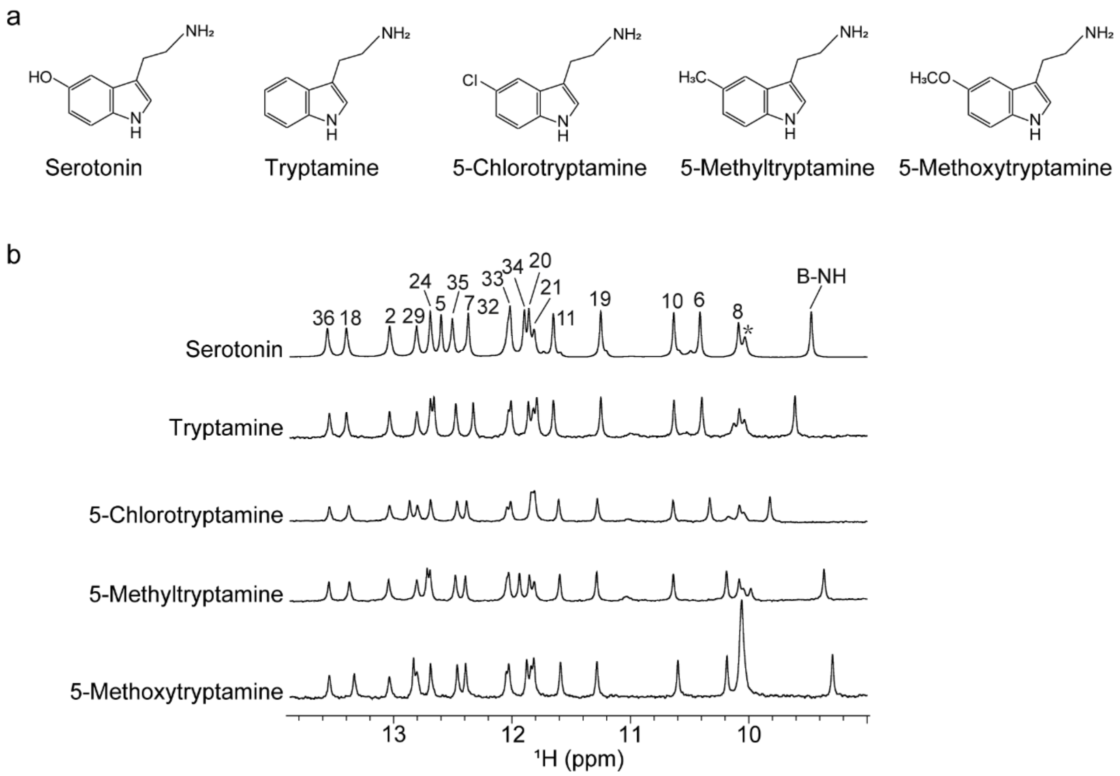


**Figure S8.** (a) Chemical structures of serotonin and its analogs. (b) Imino region of $^{1}H$ NMR spectra of Apt38 in presence of serotonin and its analogs under $K^+$ buffer conditions at 298 K. The asterisk (*) denotes the resonance from free serotonin.

**Table S1.** Proton chemical shifts of the Apt38–serotonin complex. The chemical shift values for carbon-bound (non-exchangeable) and exchangeable (active) hydrogens were derived from NMR spectra acquired in $D_2O$ and $H_2O$ buffers, respectively, at 298 K.

| Residue | H1/H3 | H41/H21 /H61 | H42/H22 /H62 | H5/Me /H2 | H1’ | H2’ | H2” | H3’ | H4’ | H5’ | H5” | H8/H6 |
|---|---|---|---|---|---|---|---|---|---|---|---|---|
| C1 | | | | 5.89 | 5.70 | 1.78 | 2.29 | 4.65 | 4.03 | 3.68 | 3.65 | 7.58 |
| G2 | 13.03 | | | | 5.24 | 2.65 | 2.68 | 4.90 | 4.22 | 4.00 | 3.90 | 7.93 |
| A3 | | | | 7.94 | 6.23 | 2.63 | 2.84 | 4.99 | 4.40 | 4.13 | 4.05 | 8.15 |
| C4 | | 8.25 | 7.11 | 5.15 | 5.76 | 1.33 | 1.97 | 4.61 | 3.89 | 4.26 | 4.02 | 6.95 |
| T5 | 12.63 | | | 1.12 | 5.01 | 2.02 | 2.19 | 4.80 | 4.11 | 3.85 | 3.91 | 6.60 |
| G6 | 10.42 | | | | 5.80 | 3.19 | 2.51 | 4.83 | 4.52 | 4.33 | 4.33 | 6.97 |
| G7 | 12.37 | | | | 5.72 | 2.89 | 2.17 | 4.97 | 4.38 | 4.14 | 4.14 | 8.39 |
| T8 | 10.09 | | | 1.44 | 5.42 | 2.27 | 2.59 | 4.97 | 4.38 | 4.01 | 3.35 | 6.10 |
| A9 | | | | 7.23 | 5.89 | 2.15 | 2.84 | 5.02 | 4.24 | 3.85 | 4.14 | 7.15 |
| G10 | 10.63 | | | | 6.04 | 3.03 | 2.87 | 4.84 | 4.47 | 4.42 | 4.22 | 7.24 |
| G11 | 11.65 | | | | 5.61 | 2.65 | 2.41 | 4.97 | 4.36 | 4.28 | 4.21 | 7.95 |
| C12 | | 8.33 | 6.91 | 5.75 | 6.64 | 1.20 | 2.07 | 4.92 | 4.31 | 4.13 | 4.13 | 7.46 |
| A13 | | | | 7.79 | 6.13 | 2.56 | 2.83 | 4.92 | 3.32 | 3.57 | 3.44 | 8.38 |
| G14 | | | | | 6.00 | 1.11 | 2.36 | 4.99 | 4.17 | | | 7.66 |
| A15 | | | | 7.87 | 6.84 | 2.66 | 3.48 | 5.44 | 4.78 | 4.35 | 4.35 | 8.55 |
| T16 | | | | 1.34 | 6.19 | 2.04 | 2.20 | 4.87 | 3.32 | 3.15 | 3.89 | 7.37 |
| A17 | | | | 8.15 | 4.94 | 2.58 | 2.73 | 4.99 | 4.33 | 3.63 | 3.44 | 7.83 |
| G18 | 13.40 | | | | 5.92 | 2.62 | 3.07 | 5.00 | 4.34 | 4.25 | 4.04 | 7.65 |
| G19 | 11.26 | 8.51 | 6.91 | | 6.59 | 2.39 | 2.06 | 4.55 | 3.86 | 4.18 | 3.89 | 7.86 |
| G20 | 11.86 | | | | 5.60 | 1.80 | 1.89 | 4.76 | 4.34 | 3.98 | 3.54 | 7.14 |
| G21 | 11.84 | | | | 4.86 | 0.80 | 1.40 | 4.66 | 3.63 | 4.24 | 4.03 | 7.46 |
| A22 | | | | 7.69 | 5.21 | 2.02 | 2.28 | 4.80 | 3.77 | 3.47 | 3.20 | 7.81 |
| A23 | | | | 8.02 | 6.04 | 1.21 | 2.28 | 4.74 | 4.35 | 3.79 | 4.04 | 7.50 |
| G24 | 12.69 | | | | 5.38 | 2.70 | 2.48 | 4.92 | 4.37 | 4.05 | 4.02 | 7.95 |
| C25 | | 6.43 | 7.94 | 5.38 | 6.11 | 2.11 | 2.22 | 4.82 | 4.27 | 4.19 | 4.09 | 7.43 |
| T26 | | | | 1.89 | 6.21 | 2.02 | 2.29 | 4.72 | 4.28 | 4.15 | 4.03 | 7.71 |
| T27 | | | | 1.66 | 5.69 | 1.93 | 2.19 | 4.58 | 3.83 | 3.93 | 3.86 | 7.51 |
| T28 | | | | 1.61 | 5.87 | 2.05 | 2.23 | 4.63 | 3.76 | 3.69 | 3.61 | 7.38 |
| G29 | 12.80 | | | | 5.93 | 2.61 | 2.78 | 4.81 | 4.32 | 3.99 | 3.93 | 7.89 |
| C30 | | 6.46 | 8.14 | 4.89 | 5.88 | 1.49 | 2.30 | 4.76 | 4.16 | 4.12 | 4.07 | 6.86 |
| G31 | | | | | 5.74 | 2.63 | 2.97 | 5.14 | 4.14 | 4.12 | 4.07 | 7.91 |
| T32 | 12.04 | | | 1.49 | 4.75 | 1.44 | 2.07 | 4.81 | | | | 7.17 |
| G33 | 12.02 | 8.01 | 5.78 | | 6.15 | 3.59 | 2.89 | 4.94 | 4.41 | 5.13 | 4.03 | 7.28 |
| G34 | 11.90 | | | | 6.62 | 3.00 | 2.87 | 5.20 | 4.68 | 4.31 | 4.36 | 8.60 |
| G35 | 12.53 | | | | 6.00 | 2.61 | 2.83 | 5.07 | 4.61 | 4.30 | 4.47 | 7.97 |
| T36 | 13.58 | | | 1.28 | 6.10 | 2.09 | 2.45 | 4.92 | 4.27 | 4.20 | 4.31 | 7.25 |
| C37 | | 8.61 | 7.06 | 5.74 | 5.64 | 2.07 | 2.35 | 4.85 | 4.13 | 4.06 | 4.09 | 7.52 |
| G38 | | | | | 6.16 | 2.64 | 2.38 | 4.69 | 4.18 | 4.06 | 4.09 | 7.96 |

**Table S2.** Proton chemical shifts of the free Apt38 complex. The chemical shift values for carbon-bound (non-exchangeable) and exchangeable (active) hydrogens were derived from NMR spectra acquired in $D_2O$ and $H_2O$ buffers, respectively, at 298 K.

| Residue | H1/H3 | H41/H21 /H61 | H42/H22 /H62 | H5/Me /H2 | H1' | H2' | H2" | H3' | H4' | H5' | H5" | H8/H6 |
|---|---|---|---|---|---|---|---|---|---|---|---|---|
| C1 | | | | 5.88 | 5.70 | 1.77 | 2.28 | 4.64 | 4.02 | 3.65 | 3.65 | 7.57 |
| G2 | 13.00 | | | | 5.26 | 2.63 | 2.66 | 4.90 | 4.22 | 4.00 | 3.89 | 7.91 |
| A3 | | | | 7.96 | 6.24 | 2.65 | 2.85 | 4.99 | 4.40 | 4.12 | 4.05 | 8.15 |
| C4 | | 8.10 | 6.91 | 5.23 | 5.90 | 1.85 | 2.19 | 4.75 | 4.04 | 4.26 | 4.11 | 7.17 |
| T5 | | | | 1.61 | 5.95 | 2.20 | 2.53 | 4.75 | 4.07 | 3.99 | 4.14 | 7.34 |
| G6 | 11.57 | | | | 5.69 | 2.83 | 2.37 | 4.88 | 4.26 | 4.18 | 4.18 | 7.20 |
| G7 | 12.20 | | | | 5.80 | 2.91 | 2.27 | 4.97 | 4.38 | | | 8.48 |
| T8 | 10.05 | | | 1.43 | 5.39 | 2.19 | 2.59 | 4.95 | 4.37 | 4.02 | 3.39 | 6.10 |
| A9 | | | | 7.32 | 5.93 | 2.26 | 2.89 | 5.04 | 4.24 | 3.87 | 4.12 | 7.20 |
| G10 | 10.71 | | | | 6.05 | 3.09 | 2.85 | 4.87 | 4.47 | 4.41 | 4.24 | 7.28 |
| G11 | 11.90 | | | | 5.60 | 2.64 | 2.36 | 4.95 | 4.37 | 4.29 | 4.20 | 7.97 |
| C12 | | 8.24 | 6.41 | 5.72 | 6.63 | 1.30 | 2.16 | 4.92 | 4.32 | 4.14 | 4.14 | 7.45 |
| A13 | | | | 7.68 | 6.11 | 2.53 | 2.80 | 4.91 | 3.34 | 3.58 | 3.44 | 8.38 |
| G14 | | | | | 5.98 | 1.18 | 2.34 | 4.97 | 4.14 | | | 7.66 |
| A15 | | | | | 6.75 | 2.65 | 3.46 | 5.40 | 4.78 | 4.32 | 4.32 | 8.52 |
| T16 | | | | 1.26 | 6.12 | 2.03 | 2.17 | 4.75 | 3.29 | 3.15 | 3.87 | 7.31 |
| A17 | | | | 7.76 | 4.90 | 2.36 | 2.69 | 4.46 | 4.40 | 3.59 | 3.33 | 7.74 |
| G18 | 13.27 | | | | 5.93 | 2.58 | 2.89 | 4.93 | 4.46 | | | 7.68 |
| G19 | 11.16 | 8.64 | 6.91 | | 6.62 | 2.42 | 2.04 | 4.53 | 3.96 | 4.07 | 3.86 | 7.84 |
| G20 | 11.78 | | | | 5.64 | 1.76 | 1.81 | 4.76 | 4.35 | 3.97 | 3.57 | 7.19 |
| G21 | 11.97 | | | | 4.91 | 0.76 | 1.44 | 4.69 | 3.70 | 4.27 | 4.05 | 7.48 |
| A22 | | | | 7.66 | 5.27 | 2.04 | 2.30 | 4.83 | 3.80 | 3.51 | 3.28 | 7.84 |
| A23 | | | | 8.02 | 6.03 | 1.13 | 2.27 | 4.73 | 4.37 | 3.81 | 4.02 | 7.50 |
| G24 | 12.69 | | | | 5.39 | 2.70 | 2.49 | 4.93 | 4.38 | 4.27 | 4.04 | 7.95 |
| C25 | | 6.44 | 7.96 | 5.38 | 6.12 | 2.10 | 2.21 | 4.82 | 4.27 | 4.18 | 4.09 | 7.43 |
| T26 | | | | 1.89 | 6.21 | 2.02 | 2.29 | 4.72 | 4.28 | 4.15 | 4.03 | 7.71 |
| T27 | | | | 1.66 | 5.70 | 1.93 | 2.19 | 4.58 | 3.83 | 3.94 | 3.86 | 7.51 |
| T28 | | | | 1.61 | 5.87 | 2.05 | 2.25 | 4.63 | 3.76 | 3.69 | 3.62 | 7.38 |
| G29 | 12.81 | | | | 5.94 | 2.60 | 2.78 | 4.81 | 4.31 | 3.99 | 3.93 | 7.89 |
| C30 | | 6.46 | 8.14 | 4.89 | 5.88 | 1.45 | 2.28 | 4.75 | 4.16 | 4.12 | 4.07 | 6.86 |
| G31 | | | | | 5.73 | 2.64 | 2.93 | 5.13 | 4.15 | 4.11 | 4.06 | 7.91 |
| T32 | 12.11 | | | 1.49 | 4.70 | 2.02 | 2.02 | 4.78 | | | | 7.15 |
| G33 | 11.98 | | | | 6.14 | 3.55 | 2.87 | 5.12 | 4.59 | | | 7.25 |
| G34 | 11.42 | | | | 6.20 | 2.85 | 2.85 | 5.12 | 4.39 | 4.28 | 4.41 | 8.48 |
| G35 | 12.31 | | | | 6.08 | 2.77 | 2.92 | 5.11 | 4.61 | 4.27 | 4.38 | 8.06 |
| T36 | 13.56 | | | 1.39 | 6.12 | 2.14 | 2.44 | 4.93 | 4.28 | 4.20 | 4.28 | 7.31 |
| C37 | | 8.58 | 7.08 | 5.76 | 5.66 | 2.08 | 2.37 | 4.85 | 4.16 | | | 7.54 |
| G38 | | | | | 6.18 | 2.64 | 2.39 | 4.69 | 4.19 | 4.08 | 4.08 | 7.97 |

**Table S3**. Intermolecular NOEs between serotonin and Apt38 aptamer protons in Apt38-serotonin complex.

| **Serotonin protons** | **Apt38 protons** |
|---|---|
| Hα | A17-H8<br>T5-H3<br>G34-H1<br>G19-H1 |
| Hβ | A17-H8<br>G19-H1<br>G34-H1 |
| Hγ | G34-H1<br>G19-H1<br>G6-H1<br>G11-H1 |
| H1 | T5-H1', H2'/H2'', H6, Me,<br>H2'/H2'', H1'<br>G6-H1<br>C12-H42 |
| H2 | T5-Me<br>G6-H1<br>C12-H41/H42 |
| H4 | G34-H1<br>T5-H3 |
| H6 | T5-H1', H2'/2''<br>G6- H4', H5'/H5'' |
| H7 | G6-H5'/H5''<br>T5-H1', H2'/H2'', H3', H6 |

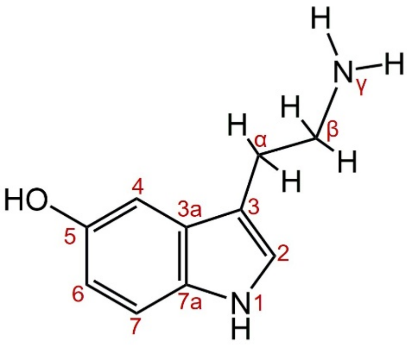

serotonin structure

**Table S4**. Statistics of the computed ten structures of free and serotonin-bound Apt38.

| | Apt38-serotonin complex | Free Apt38 |
|---|---|---|
| **Distance restraints** | | |
| Intraresidue | 65 | 64 |
| Sequencial | 164 | 167 |
| Long-range | 73 | 74 |
| Intermolecular | 39 | - |
| **Other restraints** | | |
| Hydrogen bond restraints | 109 | 104 |
| Dihedral angles | 38 | 38 |
| Plane | 13 | 12 |
| **NOE violations** | | |
| Number (>0.2 Å) | 0 | 0 |
| RMSD of vilations (Å) | 0.061 ± 0.004 | 0.058 ± 0.004 |
| **Dihedral violations** | | |
| Number (>5 $^{O}$) | 0 | 0 |
| RMSD of vilations (Å) | 0.527 ± 0.182 | 0.759 ± 0.181 |
| **Deviations from the ideal covalent geometry** | | |
| Bond lengths (Å) | 0.002 ± 0.000 | 0.002 ± 0.000 |
| Bond angles (deg) | 0.506 ± 0.012 | 0.480 ± 0.016 |
| Impropers (deg) | 0.306 ± 0.008 | 0.279 ± 0.005 |
| **Pairwise all heavy atoms RMSD values (Å)** | | |
| Entire complex | 2.64 ± 0.57 | - |
| Only Apt38 | 2.66 ± 0.58 | 2.54 ± 0.45 |
| Apt38 except residues14-16, 26-28 in loops | 1.47 ± 0.47 | 1.34 ± 0.26 |

**Table S5**. The sequence of aptamer mutants.

| Name | Sequence (5' to 3') |
|---|---|
| Apt44 | CGA CTG GTA GGC AGA TAG GGG AAG CTG ATT CGA TGC GTG GGT CG |
| Apt44-T5A | CGA CAG GTA GGC AGA TAG GGG AAG CTG ATT CGA TGC GTG GGT CG |
| Apt44-T5G | CGA CGG GTA GGC AGA TAG GGG AAG CTG ATT CGA TGC GTG GGT CG |
| Apt44-T5C | CGA CCG GTA GGC AGA TAG GGG AAG CTG ATT CGA TGC GTG GGT CG |
| Apt44-T8C | CGA CTG GCA GGC AGA TAG GGG AAG CTG ATT CGA TGC GTG GGT CG |
| Apt44-A9T | CGA CTG GTT GGC AGA TAG GGG AAG CTG ATT CGA TGC GTG GGT CG |
| Apt44-C12T | CGA CTG GTA GGT AGA TAG GGG AAG CTG ATT CGA TGC GTG GGT CG |
| Apt44-A13T | CGA CTG GTA GGC TGA TAG GGG AAG CTG ATT CGA TGC GTG GGT CG |
| Apt44-G14A | CGA CTG GTA GGC AAA TAG GGG AAG CTG ATT CGA TGC GTG GGT CG |
| Apt44-A15T | CGA CTG GTA GGC AGT TAG GGG AAG CTG ATT CGA TGC GTG GGT CG |
| Apt44-T16C | CGA CTG GTA GGC AGA CAG GGG AAG CTG ATT CGA TGC GTG GGT CG |
| Apt44-A17T | CGA CTG GTA GGC AGA TTG GGG AAG CTG ATT CGA TGC GTG GGT CG |
| Apt44-G18I | CGA CTG GTA GGC AGA TAI GGG AAG CTG ATT CGA TGC GTG GGT CG |
| Apt44-G18A | CGA CTG GTA GGC AGA TAA GGG AAG CTG ATT CGA TGC GTG GGT CG |
| Apt44-G21A | CGA CTG GTA GGC AGA TAG GGA AAG CTG ATT CGA TGC GTG GGT CG |
| Apt44-A22T | CGA CTG GTA GGC AGA TAG GGG TAG CTG ATT CGA TGC GTG GGT CG |
| Apt44-A23T | CGA CTG GTA GGC AGA TAG GGG ATG CTG ATT CGA TGC GTG GGT CG |
| Apt44-A23C | CGA CTG GTA GGC AGA TAG GGG ACG CTG ATT CGA TGC GTG GGT CG |
| Apt44-G24I | CGA CTG GTA GGC AGA TAG GGG AAI CTG ATT CGA TGC GTG GGT CG |
| Apt44-G24A | CGA CTG GTA GGC AGA TAG GGG AAA CTG ATT CGA TGC GTG GGT CG |
| Apt44-C25T | CGA CTG GTA GGC AGA TAG GGG AAG TTG ATT CGA TGC GTG GGT CG |
| Apt44-T26C | CGA CTG GTA GGC AGA TAG GGG AAG CCG ATT CGA TGC GTG GGT CG |
| Apt44-G27A | CGA CTG GTA GGC AGA TAG GGG AAG CTA ATT CGA TGC GTG GGT CG |
| Apt44-A28T | CGA CTG GTA GGC AGA TAG GGG AAG CTG TTT CGA TGC GTG GGT CG |
| Apt44-T29G | CGA CTG GTA GGC AGA TAG GGG AAG CTG AGT CGA TGC GTG GGT CG |
| Apt44-T30G | CGA CTG GTA GGC AGA TAG GGG AAG CTG ATG CGA TGC GTG GGT CG |
| Apt44-C31T | CGA CTG GTA GGC AGA TAG GGG AAG CTG ATT TGA TGC GTG GGT CG |
| Apt44-G32A | CGA CTG GTA GGC AGA TAG GGG AAG CTG ATT CAA TGC GTG GGT CG |
| Apt44-A33T | CGA CTG GTA GGC AGA TAG GGG AAG CTG ATT CGT TGC GTG GGT CG |
| Apt44-T34C | CGA CTG GTA GGC AGA TAG GGG AAG CTG ATT CGA CGC GTG GGT CG |
| Apt44-G35A | CGA CTG GTA GGC AGA TAG GGG AAG CTG ATT CGA TAC GTG GGT CG |
| Apt44-C36T | CGA CTG GTA GGC AGA TAG GGG AAG CTG ATT CGA TGT GTG GGT CG |
| Apt44-G37A | CGA CTG GTA GGC AGA TAG GGG AAG CTG ATT CGA TGC ATG GGT CG |
| Apt44-G37T | CGA CTG GTA GGC AGA TAG GGG AAG CTG ATT CGA TGC TTG GGT CG |
| Apt44-T38C | CGA CTG GTA GGC AGA TAG GGG AAG CTG ATT CGA TGC GCG GGT CG |
| Apt44-G4C41 | CGA GTG GTA GGC AGA TAG GGG AAG CTG ATT CGA TGC GTG GCT CG |
| Apt44-T4A41 | CGA TTG GTA GGC AGA TAG GGG AAG CTG ATT CGA TGC GTG GAT CG |
| Apt44-A4T41 | CGA ATG GTA GGC AGA TAG GGG AAG CTG ATT CGA TGC GTG GTT CG |
| Apt44-add1bp | G CGA CTG GTA GGC AGA TAG GGG AAG CTG ATT CGA TGC GTG GGT CGC |
| Apt44-add2bp | CG CGA CTG GTA GGC AGA TAG GGG AAG CTG ATT CGA TGC GTG GGT CGC G |
| Apt44-cut1bp | GA CTG GTA GGC AGA TAG GGG AAG CTG ATT CGA TGC GTG GGT C |
| Apt44-cut2bp | A CTG GTA GGC AGA TAG GGG AAG CTG ATT CGA TGC GTG GGT |
| Apt44-cut3bp | CTG GTA GGC AGA TAG GGG AAG CTG ATT CGA TGC GTG GG |

**Table S6**. The sequence of aptamer mutants.

| Name | Sequence (5' to 3') |
|---|---|
| Apt44 | CGA CTG GTA GGC AGA TAG GGG AAG CTG ATT CGA TGC GTG GGT CG |
| Apt44-loopCut1(Apt39) | CGA CTG GTA GGC AGA TAG GGG AAG CT TT TGC GTG GGT CG |
| Apt44-loopCut2(Apt38) | CGA CTG GTA GGC AGA TAG GGG AAG CT TT GC GTG GGT CG |
| Apt44-loopCut3(Apt37) | CGA CTG GTA GGC AGA TAG GGG AAG CT T GC GTG GGT CG |
| Apt44-loopCut4(Apt36-1) | CGA CTG GTA GGC AGA TAG GGG AAG CT GC GTG GGT CG |
| Apt44-loopCut5(Apt36-2) | CGA CTG GTA GGC AGA TAG GGG AAG T TT C GTG GGT CG |
| Apt44-loopCut6(Apt35) | CGA CTG GTA GGC AGA TAG GGG AA T TT T GTG GGT CG |
| Apt38-T8A | CGA CTG GAA GGC AGA TAG GGG AAG CT TT GC GTG GGT CG |
| Apt38-A9C | CGA CTG GTC GGC AGA TAG GGG AAG CT TT GC GTG GGT CG |
| Apt38-G21C | CGA CTG GTA GGC AGA TAG GGC AAG CT TT GC GTG GGT CG |

**Table S7**. ITC-determined binding affinities of aptamer mutants.

| Name | N (sites) | $K_d$ (nM) | ΔH (kcal/mol) | ΔG (kcal/mol) | -TΔS (kcal/mol) |
|---|---|---|---|---|---|
| Apt44 | 0.591 ±0.003 | 37.0 ± 4.3 | -21.3 ± 0.192 | -10.1 | 11.1 |
| Apt44-T5A | 0.528 ±0.052 | 3510 ± 1290 | -20.1 ± 4.65 | -7.44 | 12.7 |
| Apt44-T5G | | NB | | | |
| Apt44-T5C | | NB | | | |
| Apt44-T8C | | NB | | | |
| Apt44-A9T | | NB | | | |
| Apt44-C12T | | NB | | | |
| Apt44-A13T | 0.845 ±0.005 | 49.1 ± 6.8 | -25.0 ± 0.304 | -9.98 | 15.1 |
| Apt44-G14A | 0.401 ± 0.004 | 131 ± 16 | -23.6 ± 0.414 | -9.40 | 14.2 |
| Apt44-A15T | 0.555 ±0.004 | 184 ± 14 | -26.0 ± 0.294 | -9.19 | 16.9 |
| Apt44-T16C | 0.484 ±0.004 | 39.7± 7.6 | -21.1 ± 0.320 | -10.1 | 11.0 |
| Apt44-A17T | | NB | | | |
| Apt44-G18I | | NB | | | |
| Apt44-G18A | | NB | | | |
| Apt44-G21A | | NB | | | |
| Apt44-A22T | | NB | | | |
| Apt44-A23T | | NB | | | |
| Apt44-A23C | 0.190 ±0.012 | 625 ± 199 | -35.2 ± 4.88 | -8.47 | 26.7 |
| Apt44-G24I | | NB | | | |
| Apt44-G24A | | NB | | | |
| Apt44-C25T | 0.221 ±0.003 | 28.8± 5.6 | -21.4 ± 0.426 | -10.3 | 11.1 |
| Apt44-T26C | 0.334 ±0.003 | 24.6 ± 5.0 | -20.6 ± 0.320 | -10.4 | 10.3 |
| Apt44-G27A | 1.14 ± 0.012 | 55.6 ± 14.6 | -22.1 ± 0.587 | -9.90 | 12.2 |
| Apt44-A28T | 0.695 ±0.003 | 27.4 ± 3.5 | -22.4 ± 0.197 | -10.3 | 12.1 |
| Apt44-T29G | 0.166 ±0.002 | 33.8 ± 9.6 | -23.0 ± 0.534 | -10.2 | 12.8 |
| Apt44-T30G | 0.102 ±0.005 | 14.7 ± 10.5 | -22.2 ± 1.62 | -10.7 | 11.5 |
| Apt44-C31T | 0.766 ±0.005 | 45.5 ± 8.0 | -21.4 ± 0.319 | -10.0 | 11.4 |
| Apt44-G32A | 0.857 ±0.004 | 31.4 ± 4.8 | -21.6 ± 0.248 | -10.2 | 11.3 |
| Apt44-A33T | 0.614 ±0.002 | 42.7 ± 3.8 | -22.0 ± 0.164 | -10.1 | 12.0 |
| Apt44-T34C | 0.212 ±0.002 | 39.1 ± 5.5 | -22.8 ± 0.365 | -10.1 | 12.7 |
| Apt44-G35A | | NB | | | |
| Apt44-C36T | | NB | | | |
| Apt44-G37A | | NB | | | |
| Apt44-G37T | | NB | | | |
| Apt44-T38C | | NB | | | |
| Apt44-G4C41 | 0.115 ±0.003 | 321 ± 156 | -23.8 ± 7.02 | -8.86 | 14.9 |
| Apt44-T4A41 | 0.241 ±0.032 | 1750 ± 426 | -36.5 ± 7.14 | -7.85 | 28.7 |
| Apt44-A4T41 | 0.359 ±0.014 | 415 ± 104 | -26.4 ± 1.69 | -8.71 | 17.7 |
| Apt44-add1bp | 0.266 ±0.002 | 33.2 ± 6.8 | -21.0 ± 0.331 | -10.2 | 10.8 |

| | | | | | |
|---|---|---|---|---|---|
| Apt44-add2bp | 0.202 ±0.003 | 29.5 ± 5.5 | -20.6 ± 0.386 | -10.3 | 10.3 |
| Apt44-cut1bp | 0.719 ±0.004 | 134 ± 10 | -32.0 ± 0.307 | -9.38 | 22.6 |
| Apt44-cut2bp | 0.822 ±0.176 | 10400 ± 11000 | -41.8 ± 35.6 | -6.80 | 35.0 |
| Apt44-cut3bp | | NB | | | |
| Apt44-loopCut1(Apt39) | 0.902 ±0.004 | 29.6 ± 4.7 | -21.6 ± 0.253 | -10.3 | 11.3 |
| Apt44-loopCut2(Apt38) | 0.910 ±0.002 | 25.4 ± 2.1 | -21.7 ± 0.127 | -10.4 | 11.3 |
| Apt44-loopCut3(Apt37) | 1.04 ±0.004 | 29.9 ± 3.5 | -21.6 ± 0.199 | -10.3 | 11.3 |
| Apt44-loopCut4(Apt36-1) | 0.185 ±0.002 | 24.2 ± 6.5 | -23.3 ± 0.450 | -10.4 | 12.9 |
| Apt44-loopCut5(Apt36-2) | 0.958 ±0.002 | 19.7 ± 1.9 | -21.3 ± 0.138 | -10.5 | 10.8 |
| Apt44-loopCut6(Apt35) | 0.381 ±0.004 | 116 ± 15 | -28.3 ± 0.483 | -9.47 | 18.8 |
| Apt38-T8A | | NB | | | |
| Apt38-A9C | | NB | | | |
| Apt38-G21C | | NB | | | |

Binding buffer: 1×PBS, 2 mM $MgCl_2$, 298K

NB: no binding

**Table S8**. ITC-determined binding affinities of Apt44 for structural analogs of serotonin.

| **Analogs** | **N (sites)** | **$K_d$ (nM)** | **ΔH (kcal/mol)** | **ΔG (kcal/mol)** | **TΔS (kcal/mol)** |
|---|---|---|---|---|---|
| Tryptophol | | NB | | | |
| L-tryptophan | | NB | | | |
| 5-Hydroxy-L-tryptophan | | NB | | | |
| Tryptamine | 0.706 ± 0.005 | 12.9 ± 2.8 | -24.5 ± 0.372 | -10.8 | 13.7 |
| 5-Chlorotryptamine | 0.441 ± 0.005 | 3.6 ± 2.4 | -25.3 ± 0.474 | -11.5 | 13.8 |
| 5-Methyltryptamine | 0.368 ± 0.005 | 3.2 ± 3.9 | -30.1 ± 0.884 | -11.6 | 18.5 |
| 5-Methoxyltryptamine | 0.489 ± 0.004 | 5.6 ± 1.9 | -29.0 ± 0.445 | -11.3 | 17.8 |

Binding Buffer: 1×PBS (pH 7.5), 2 mM $MgCl_2$, 298 K

NB: no binding
